# The mitochondrial methylation potential gates mitoribosome assembly

**DOI:** 10.1101/2024.11.15.623401

**Authors:** Ruth I.C. Glasgow, Vivek Singh, Lucía Peña-Pérez, Alissa Wilhalm, Marco F. Moedas, David Moore, Florian A. Rosenberger, Xinping Li, Ilian Atanassov, Mira Saba, Miriam Cipullo, Joanna Rorbach, Anna Wedell, Christoph Freyer, Alexey Amunts, Anna Wredenberg

## Abstract

S-adenosylmethionine (SAM) is crucial for cellular processes, primarily serving as the principal methyl group donor of the cell and playing a key role in gene regulation and translation on the ribosome. Inside mitochondria, SAM-dependent methylations occur at several steps of gene expression, but their role and significance remain unclear. Using direct long-read RNA sequencing on mouse tissue and mouse embryonic fibroblasts, we demonstrate that the mitochondrial ribosomal gene cluster is not efficiently processed without mitochondrial SAM. This results in the accumulation of unprocessed ribosomal RNA precursors. Protein profiling of ribosome fractions revealed that these precursors are associated with processing and ribosome assembly factors, indicating stalling at an early stage. Structural analysis of the mitochondrial ribosome by cryogenic electron microscopy revealed that mitochondrial SAM is required during peptidyl transferase centre formation and mitochondrial ribosome assembly. Our data thus identify a critical role for methylation at two steps during mitochondrial gene expression.

## Introduction

S-adenosylmethionine (SAM) is a product of one-carbon metabolism and the primary biological methyl group donor within our cells^1^. Its levels are implicated in numerous cellular processes, including the cell’s epigenetic state or as a substrate for the biosynthesis of various macromolecules. SAM is synthesised in the cytosol from adenosine 5ʹ-triphosphate (ATP) and methionine as part of the methionine cycle before relocating to different subcellular compartments. Approximately 30% of cellular SAM resides inside mitochondria, acting as a cofactor for ubiquinone and lipoic acid biosynthesis and as a methyl group donor for mitochondrial RNAs and various proteins^2^. Mitochondrial gene expression especially relies on mitochondrial SAM (mitoSAM), with multiple nucleotide modifications, such as base or 2ʹ-*O*-ribose methylations, reported on mitochondrial transfer and ribosomal RNAs^3,4^.

The mammalian mitochondrial genome (mtDNA) is expressed as long, polycistronic transcripts that cover almost the entire genome length. The canonical mode of mitochondrial RNA processing describes how tRNAs act as recognition sites for dedicated nucleases to sequentially release the flanking transcripts for further maturation in a 5ʹ to 3ʹ order^5^. The molecular and structural details of this process have been described, with the mitochondrial ribonucleases (mtRNase) P and Z cleaving the 5ʹ and 3ʹ tRNA junctions, respectively^6–10^. mtRNase P consists of the two tRNA-binding subunits, MRPP1 and 2, and the catalytic subunit MRPP3, also known as PRORP^10^. MRPP1, also known as RNA methyltransferase 10 homolog C, TRMT10C, additionally catalyses the formation of N(1)-methylguanine (m^1^G) and N(1)-methyladenine (m^1^A) at position 9 in 19 out of 22 mitochondrial tRNAs^3,11^. Recent structural work, though, indicates that this methylation is not required for cleavage by either RNase P or Z^9,12,13^.

At least eight residues of the mammalian small (12S) and large (16S) mitochondrial ribosomal subunit RNAs are methylated^14–16^. The responsible SAM-dependent methyltransferases have been identified, with TRMT2B, METTL15, NSUN4, and TFB1M modifying residues of 12S^17–23^, and MRM1, MRM2, and MRM3 responsible for 2ʹ-*O*-ribose methylations on 16S^24–26^. In mice, 12S rRNA only contains base methylations (m^5^U499, m^4^C909, m^5^C911, m^6^_2_A1006, m^6^_2_A1007), while 16S rRNA only has 2ʹ-*O*-ribose methylations (Gm2253, Um2481, Gm2482) (mouse mtDNA (NC_005089) numbering shown). Human 16S rRNA contains an additional methylation at m^1^A946 by TRMT61B^27^.

The roles of these modifications are not always clear, but the recent structure of a fully modified human mitoribosome shows that six mitochondria-specific modifications help stabilise the decoding centre and facilitate the binding of the mRNA and tRNA ligands^23^. For instance, cytosine N^4^- and N^5^-methylation (m^4^C and m^5^C) by METTL15 and NSUN4 on 12S rRNA have recently been implicated in monosome assembly and translation^20,22,23^. These modifications have been shown to interact with the phosphate group of m^4^C1486 (m^4^C909 in mice), thereby stabilising an mRNA kink between the A- and P-sites.^20,22,23^. Additionally, on 16S rRNA, 2ʹ-*O*-ribose methylations by MRM1-3 position the rRNA bases sterically to enhance interactions with the A-tRNA nucleotides^23^. Importantly, though, MRM2 methylation activity has recently been suggested to be dispensable for ribosome assembly^28^.

SAM cannot be *de novo* synthesised inside mitochondria and must be imported via the mitochondrial SAM carrier (SAMC), encoded by *SLC25A26*^2,29–31^. Variants in *SLC25A26* can cause severe mitochondrial disease in humans^32–34^, and the deletion of SAMC in mice and flies is embryonic and larvae lethal^35^. We previously replicated patient variants in fly models, demonstrating that a reduced SAM import into mitochondria can result in a progressive mitochondrial gene expression defect, but the mechanism remains unknown^35^.

To investigate the undefined importance of mitochondrial RNA methylation we attenuated the mitochondrial methylation potential in a conditional skeletal muscle-specific mouse model and mouse embryonic fibroblasts (MEFs). We show that efficient processing of the mitochondrial ribosome gene cluster depends on mitoSAM. We also demonstrate that the assembly of the small ribosomal subunit (SSU) begins before the complete processing of 12S rRNA. Finally, our data reveal methylation as a critical PTC assembly and the large ribosomal subunit (LSU) maturation checkpoint, highlighting a fundamental role for SAM during mitochondrial gene expression.

## Results

### Mitochondrial SAM depletion causes a progressive defect in gene expression

We previously reported that SAMC is the only route for SAM into mitochondria, and its deletion drastically depletes mitochondria of SAM^35^. While the *in vivo* deletion is lethal and presents a progressive mitoSAM deficiency, SAMC KO MEFs (KO^MEF^) can be maintained in culture and represent a chronic state of mitoSAM depletion. To study the progressive decline in mitochondrial methylation potential, we deleted *Slc25a26* specifically in mouse skeletal muscle, using *Cre* recombinase under the control of the myosin light chain 1f promoter (*Mlc1f-Cre*). Muscle-specific SAMC KO (KO^SkM^) mice were born at Mendelian ratios and appeared normal at weaning but needed to be sacrificed by 12 weeks of age due to plateaued weight. We measured m^4^C909 and m^5^C911 methylation of 12S rRNA by bisulphate sequencing as a readout of the mitochondrial methylation potential. At 8 weeks of age, quadriceps presented with less than 50% methylation compared to controls, with a further decrease by 12 weeks (Figure 1a). KO^MEF^ cells only retained residual methylation at both sites, confirming the lack of methylation potential in these cells.

**Figure 1.**
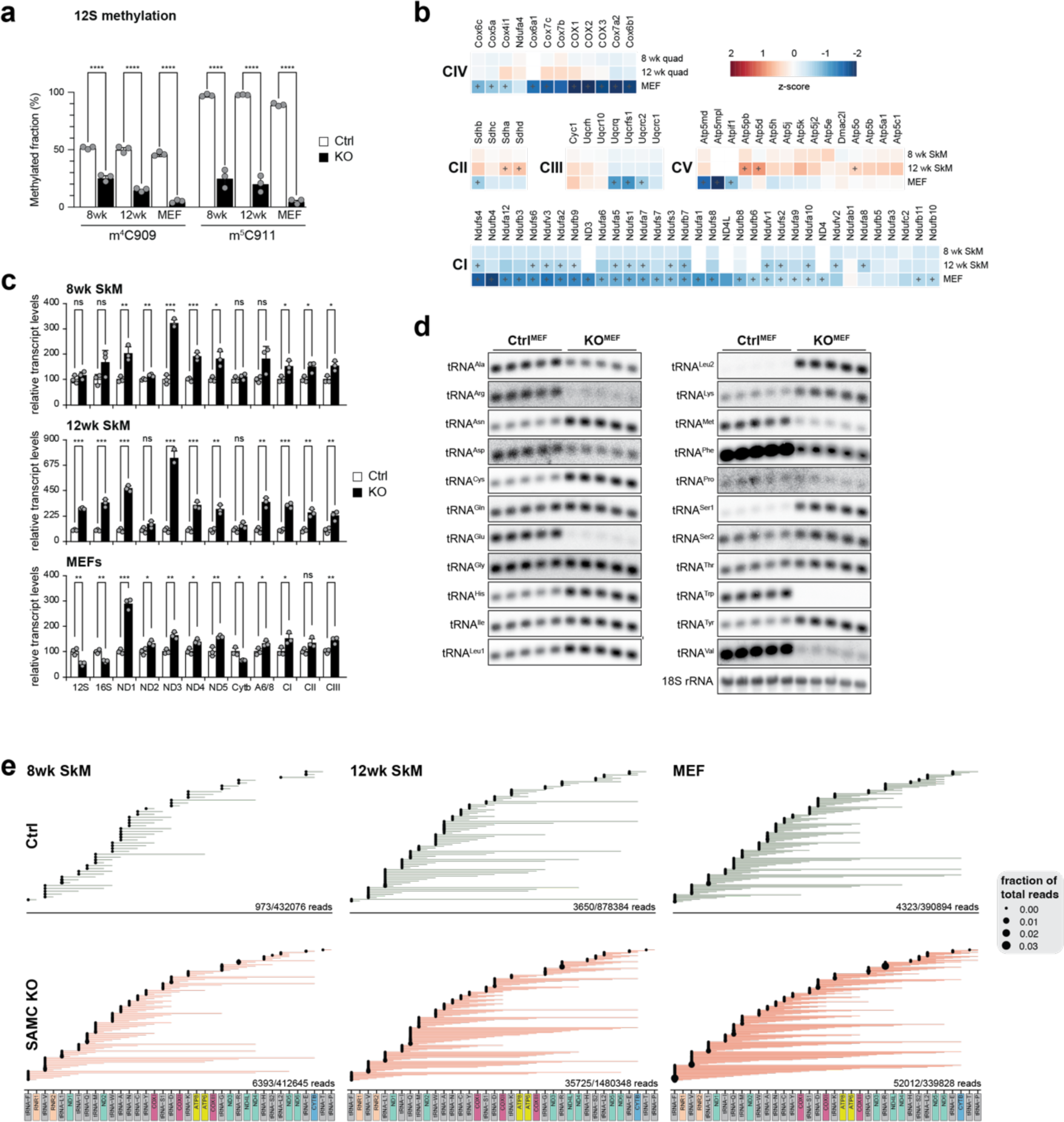
Reduced mitochondrial methylation potential affects mitochondrial gene expression. **a**, The mitochondrial methylation potential was determined by bisulfite pyrosequencing of 12S mt-rRNA from control (white) and SAMC KO (black) MEFs or quadriceps at 8 and 12-weeks of age, targeting 4ʹ-methylcytosine m^4^C909 (m^4^C) or 5ʹ-methylcytosine m^5^C911 (m^5^C). (n = 3) **b**, Heatmap of OXPHOS subunits as determined by mass spectrometry-based proteomics in 8 and 12-week-old quadriceps or MEF samples normalised to controls (n = 3). **c**, Relative mitochondrial transcript steady-state levels in 8 and 12-week-old muscle or MEFs from control (white) and SAMC KO (black) samples, as determined by qRT-PCR (n =3 biologically independent samples with 3 technical replicates). **d**, Mitochondrial tRNA steady-state levels in SAMC KO (black) or control (white) MEFs, as determined by Northern blot analysis (n =5 biologically independent samples). **e**, ONT sequencing data filtered for all reads containing tRNA sequences and mapped against the mitochondrial genome (KR020497). Control (Ctrl; green) and SAMC KO (red) from 8 and 12-week-old muscle (quad) and MEF samples (n=3 biologically independent samples). Student’s two-tailed T-test was used. Error bars represent Standard deviation. ns p > 0.05; *p < 0.05, **p < 0.01, ***p < 0.001.

Quantitative proteomics, comparing control to KO^SkM^ quadriceps at 8 and 12 weeks of age, revealed a mild but progressive decline in individual OXPHOS subunit proteins of complexes I, III, and IV, and a slight increase in complex II and V subunits with age (Figure 1b and Table S1). Conversely, KO^MEFs^ exhibit a generalised decrease in OXPHOS subunits, with complexes I and IV primarily affected^35^.

Assessment of mt-mRNA transcript levels ruled out a depleted transcript pool as the cause for the decreased OXPHOS composition as steady-state levels of most transcripts were increased in KO^SkM^ quadriceps at 8 weeks of age and further rose by 12 weeks (Figure 1c). KO^MEFs^ also exhibited elevated steady-state levels with *Nd1* most changed. Only *12S*, *16S*, and *Cytb* RNA levels decreased to ∼50% of controls^35^.

We next investigated mt-tRNA steady-state levels in quadriceps and MEFs as most mt-tRNAs are post-transcriptionally methylated, including the m^1^A methylation at position 9 by TRMT10C^3,10,11^. Northern blot analysis revealed a very similar pattern in both models, although the responses to the loss of methylation of individual mt-tRNAs varied drastically (Figure 1d and Figure S1). For instance, while the steady-state levels for *mtR*, *mtE*, *mtW*, and *mtV* were almost undetectable, *mtN*, *mtC*, *mtH*, *mtL1*, *mtL2*, *mtS1, mtK, and mtY* were increased, irrespective of their strand or genome position.

### The mitochondrial methylation potential affects the processing of primary transcripts

To test if mitoSAM is required during canonical transcript processing, we performed direct long-read RNA sequencing by Oxford nanopore technology (ONT sequencing) on polyadenylated total RNA from 8- and 12-week-old mouse quadriceps and mitochondrial RNA extracts from MEF samples (Table S2) (see materials and methods). We aligned all reads to the mitochondrial genome, resulting in relatively equal read counts within the muscle or MEF groups (Figure S2a and Table S3-9). Filtering for incompletely processed transcripts (i.e. bicistronic or polycistronic reads containing at least one mt-tRNA) revealed a stark increase of multicistronic transcripts already in 8-week-old KOSkM quadriceps, which progressed by 12 weeks and was most prominent in KOMEFs (Figure 1e). Interestingly, low levels of such polycistronic transcripts were also observed in control samples, some covering almost the entire genome. While representing a small proportion of total reads in control samples (0.2% and 1.1% of total mitochondrial reads in control quadriceps and MEFs, respectively), these processing intermediates accumulated between 8 and 12 weeks of age in quadriceps (1.5% and 2.4%, respectively) and reached up to 15% of total mitochondrial reads in SAMC KO MEF cells.

We next filtered the ONT sequencing data for reads corresponding to unprocessed transcripts representing at least 0.01% of all mitochondrial reads (Figure 2a). The number of multicistronic transcripts inversely correlated with the mitochondrial methylation potential, with KO^MEFs^ containing most unprocessed transcripts. Three genomic regions were especially prone. These were (i) both rRNAs and flanking regions; (ii) *Nd2* and the upstream/downstream tRNA clusters that could span the entire *WANCY* tRNA cluster; and (iii) *mtR* together with the *Nd4/4l* bicistronic transcript (Figure 2a). We focused on the ribosomal gene cluster as together these unprocessed species accumulated to a large proportion of the total reads. The most prevalent species consisted of *mtF*-*12S*-*mtV*-*16S-mtL1*, previously referred to as RNA4^36,37^, suggesting that processing of this transcript might be under specific control compared to the remaining genome. Notably, several tRNAs, including *mtF*, *mtL1*, and *mtI*, retained their 5ʹ-leader or 3ʹ-trailer sequences, deviating from the proposed strict 5ʹ to 3ʹ processing hierarchy^38^. However, *mtV* was never observed with only a 5ʹ-leader sequence, suggesting that 5ʹ cleavage of *mtV* is a pivotal step in the processing of the ribosomal gene cluster. We further mapped the start and end sites of all reads containing 12S, 16S or *Nd1* in KO^MEFs^ and KO^SkM^ quadriceps (Figure 2b-d and Figure S3-5). For instance, in KO^MEFs^, only around 6% of 12S and 16S, and 40% of *Nd1* transcripts, were fully processed (Figure S2b). Curiously, although ∼90% of 12S transcripts were fully released in control MEFs, the majority of 16S transcripts (85%) were incompletely processed, indicating a clear preference towards 12S rRNA processing.

**Figure 2.**
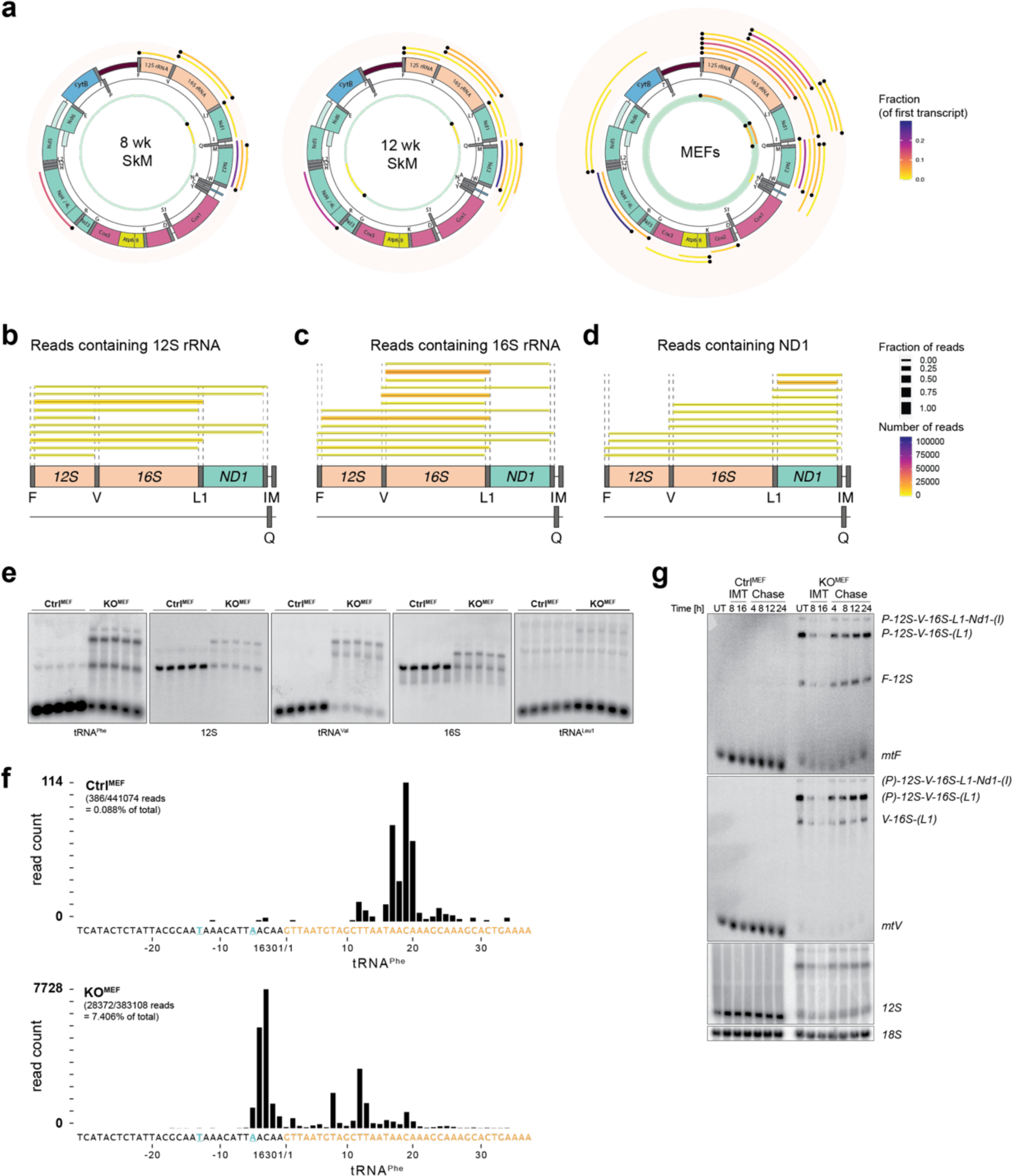
Mitochondrial SAM is critical for processing of the rRNA gene cluster. **a**, ONT sequencing data filtered for reads containing tRNA sequences present in more than 0,01% of the total RNA pool and mapped against the mitochondrial genome (KR020497). Control (inner circle; green) and SAMC KO (outer circle; pink) reads shown from 8 and 12-week-old quadriceps and MEF samples. Black dots represent presence of a tRNA sequence at the 5ʹ or 3ʹ termini of the reads. (n=3 biologically independent samples). **b**-**d**, Zoom-in of KO^MEFs^ RNA reads passing through (b) 12S rRNA, (c) 16S rRNA or (d) Nd1 with no fraction cut-off. Gene borders are indicated by dotted lines. (n=3 biologically independent samples). e, Northern blot analysis of rRNA mitochondrial transcripts and flanking tRNAs in control and SAMC KO MEFs as indicated (n =5 biologically independent samples). f, Number and position of reads containing mtF (tRNAPhe; orange) in KOMEFs is shown. Transcription start sites for the heavy strand promoter are indicated in blue. (n=3 biologically independent samples). Number of reads and percentage of total reads is indicated. **g**, Northern blot analysis of selected mitochondrial transcripts in control or SAMC KO MEFs after treatment with 20mM inhibitor of mitochondrial transcription (IMT), using probes against mtF (top panel), mtV (middle panel) or 12S (bottom panel). 18S was used as loading control. Treatment times were 8 and 16 hours, chase timepoints were at 4, 8, 12, and 24 hours. UT = untreated (dimethyl sulfoxide (DMSO) vehicle without IMT). Representative experiment of three independent experiments.

We confirmed these processing intermediates by Northern blot analysis, where the rRNA gene cluster was the predominantly affected region (Figure 2e and Figure S6). Comparing ONT sequencing coverage with qRT-PCR results highlighted an overrepresentation of 12S and 16S rRNA in KO^MEFs^ while increased *Nd1* and decreased *Cytb* steady-state levels were consistent between the two datasets (Figure 1c and Figure S2a). This discrepancy may reflect the polyA-enrichment of samples for ONT sequencing capturing rRNA attached to the more extensively polyadenylated *Nd1*.

Due to technical limitations, ONT sequencing does not sequence the 5ʹ ends of transcripts, and the reads consistently mapped ∼12nt (± 2) downstream of the reported start sites (Figure S7). In agreement, in control samples, the 5ʹ ends of transcripts containing *mtF* mapped ∼20nt downstream of its annotated start site (Figure 2f and Figure S8)^39^. In contrast, in KO^SkM^ and KO^MEF^ samples, most start sites mapped further upstream, consistent with the reported heavy strand promoter (HSP) start site, thus demonstrating failed processing of the 5ʹ untranslated region of *mtF*.

To determine the stability of these unprocessed transcripts, we treated control and KO MEFs with an inhibitor of mitochondrial transcription (IMT)^40^. While the steady-state levels of fully processed transcripts gradually decreased in both IMT-treated cell lines (Figure 2g), unprocessed intermediates exhibited a more rapid clearance, suggesting a faster turnover. Upon IMT removal, these species reappeared, implying that these rRNA processing intermediates respond dynamically to transcription rates. Thus, this data identify methylation within the rRNA gene cluster as critical for efficient processing.

### Mitochondrial ribosome assembly depends on mitochondrial SAM

To investigate whether the altered mt-rRNA and mt-tRNA compositions affected translation, we examined the mitochondrial translation capacity in mitochondria isolated from quadriceps or MEFs. Mitochondrial translation was mildly reduced in 12-week-old KO^SkM^ samples, and no signal from nascent peptides was detected in KO^MEFs^ (Figure 3a). To determine if this defect is specific to the altered mt-tRNA composition or monosome formation, we performed sedimentation experiments on the mitochondrial ribosome. KO^SkM^ samples presented a normal sedimentation pattern, suggesting that the modified tRNA composition is the leading cause of the observed translation defect in these muscle samples (Figure S9a). In contrast, chronic depletion of mitoSAM in MEFs resulted in a reduced monosome formation, with normal signals corresponding to the mtLSU and mtSSU (Figure 3b). Mass spectrometry-based label-free proteomics (Figure 3c) and Western blot analysis (Figure 3d) demonstrated that several factors of the mitochondrial translation machinery were markedly increased in KO^SkM^ quadriceps but remained unchanged or even reduced in KO^MEFs^, suggesting a compensatory response in the muscle (Figure 3c,d)^35^. Thus, while muscle can maintain nearly normal monosome formation and translation in an attempt to compensate for the OXPHOS defect, the sustained depletion of mitoSAM in MEFs affects efficient monosome formation.

**Figure 3.**
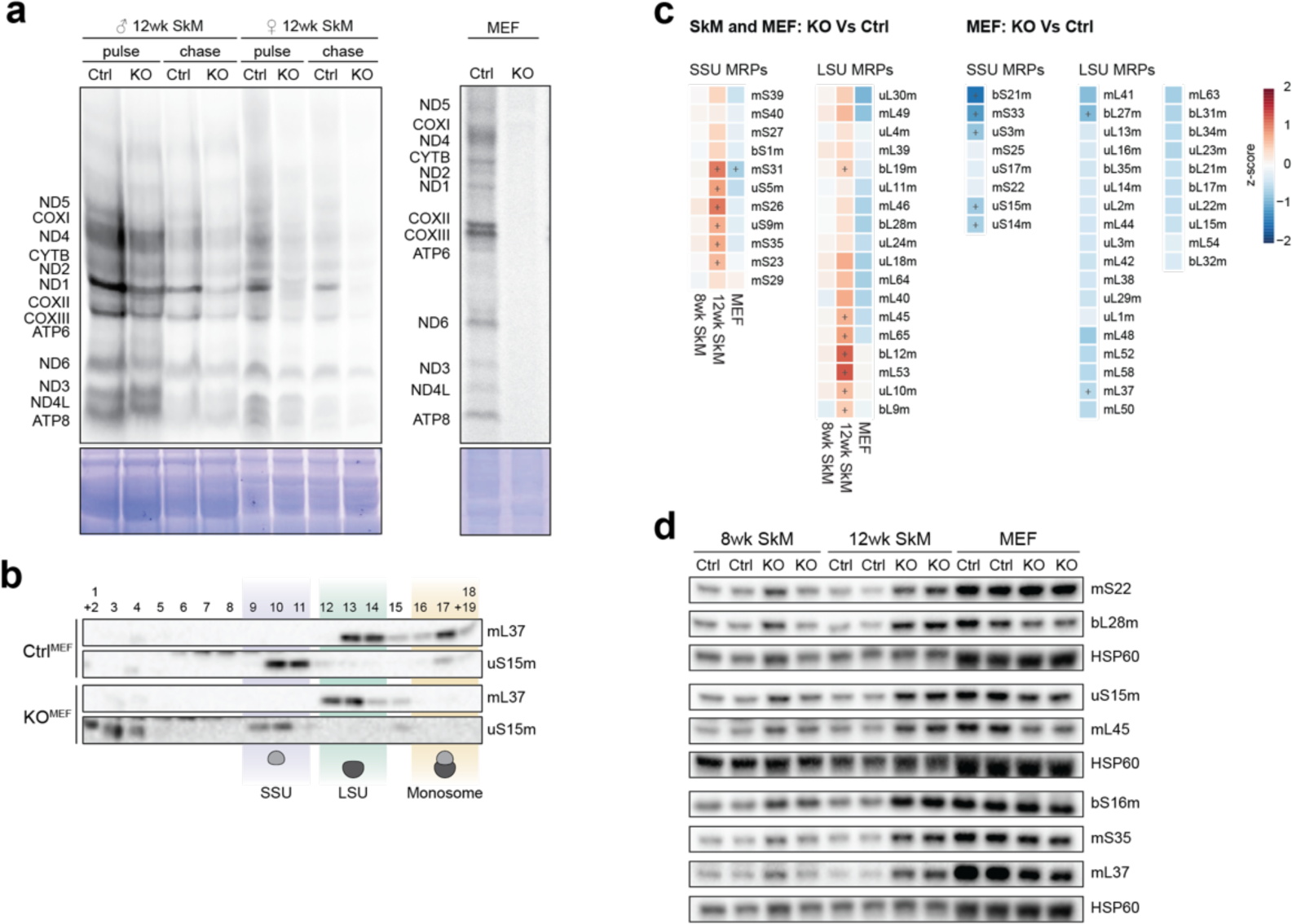
Mitochondrial ribosome assembly depends on SAM. **a**, De novo translation in isolated mitochondria from male and female 8 week and 12-week-old muscle or MEFs samples as indicated. Expected mitochondrial proteins are shown. Coomassie stain of the gel indicates loading. **b**, Western blot analysis of ribosome gradient fractions from SAMC KO and control MEFs, probed with antibodies against the small (uS15m) and large (mL37) mitochondrial ribosome subunits. The small (28S), large (39S), and monosome (55S) fractions are indicated. A representative experiment is shown of three independent experiments performed with independently prepared samples. **c**, Proteomic levels of mitoribosome subunits from 8 and 12-week-old muscle or MEF SAMC KO samples normalised to control samples. (n = 3). **d**, Western blot analysis of mitochondrial ribosome subunits in 25 ug of mitochondrial lysate from 8 and 12-week-old muscle or MEF samples.

We next asked whether immature mtSSU and mtLSU were the cause of the failed monosome formation. To investigate this, we performed stable isotope labelling with amino acids in cell culture (SILAC) on MEFs, followed by separation of mitochondrial ribosomes on a sucrose gradient and mass spectrometry of fractions corresponding to the mtSSU, mtLSU, and monosome to identify their respective compositions (Figure S9b-f and Table S10). Generally, monosome formation requires an equal stoichiometry of mtSSU and mtLSU proteins. However, although we observed an overall decrease in ribosomal proteins in these fractions in the KO sample (Figure S9e), the distribution of mtSSU and mtLSU ribosomal proteins differed greatly between the fractions. Levels of SSU proteins increased between the mtSSU and monosome fractions, whereas mtLSU proteins decreased in both the LSU and monosome fractions (Figure 4a,b). Interestingly, five mtSSU proteins did not increase in the monosome fraction, of which uS11m, uS12m, bS21m and mS33 all have recently been suggested to be ‘late-binding MRPs’^41^.

**Figure 4.**
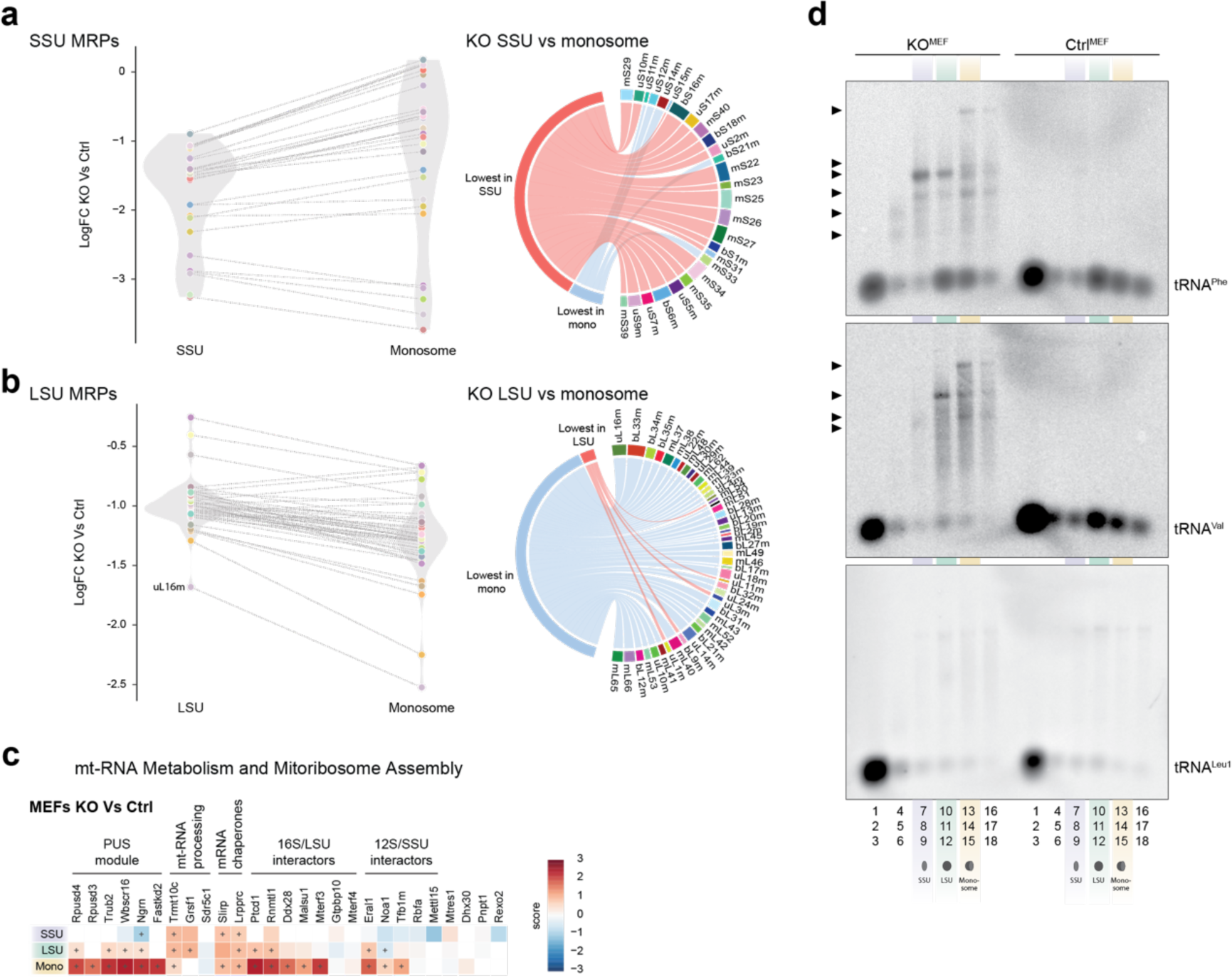
The unprocessed mitochondrial rRNA cluster engage in early ribosome assembly. **a,b**, Distribution of subunit levels of the (a) small or (b) large mitochondrial ribosome in fractions representing the mtSSU or monosome after SILAC labelling of SAMC KO and control MEFs (n=2). Right panel shows changes of individual subunits that are either increased (red) or decreased (blue) in the monosome fraction. Right panel Log fold change, and distribution of (a) mtSSU (b) mtLSU factors in the respective fractions. **c**, Enrichment levels of mitochondrial RNA binding proteins and ribosome assembly proteins in ribosome gradient fractions in SAMC KO and control MEFs after SILAC labelling (n=2). **d**, Northen blot analysis of ribosome gradient fractions from SAMC KO and control MEFs (n=1).

Surprisingly, we also observed a strong enrichment of several mitochondrial RNA binding proteins typically associated with processing and early stages of RNA maturation in the higher molecular weight fractions (15-17) of the KO^MEFs^ samples (Figure 4c). For instance, TRMT10C was enriched in all three fractions, and several factors important during the early stages of the mtSSU, such as NOA1, ERAL1, and TFB1M, as well as the entire pseudouridylation (PUS) module, which is involved in 16S rRNA modification, accumulated in the monosome fraction. We confirmed this distribution for RPUSD4 and ERAL1 on ribosome gradients using Western blot analysis (Figure S9g). Interestingly, despite exhibiting a milder translation defect and normal mitoribosome sedimentation patterns, KO^SkM^ gradients also showed an enrichment of RPUSD4 in LSU and monosome fractions (Figure S9a), demonstrating the same mechanism for rRNA processing intermediate stabilisation *in vivo*.

Taken together, these results suggest that in the absence of methylation, the mtSSU and mtLSU maturation arrests in immature states. Additionally, early processing and maturation factors in the monosome fractions indicate that a larger protein-RNA complex already assembles prior to complete cleavage of the ribosomal gene cluster.

### Processing of mt-tRNA^Phe^ and mt-tRNA^Val^ promotes early ribosome assembly

To test whether the unprocessed rRNA gene cluster is part of a larger complex, we isolated RNA from sucrose gradient fractions, followed by Northern blot analysis using probes against *mtF*, *mtV*, and *mtL1*. In control MEFs, all three processed tRNAs sedimented with the free fractions (fractions 1-3). Interestingly, though, *mtF* and *mtV*, but not *mtL1*, also co-migrated with fractions corresponding to the LSU and the monosome (fractions 12-15) (Figure 4d). The mitochondrial ribosome differs from cytosolic and bacterial ribosomes in that besides containing 12S and 16S rRNAs, it also includes a structural tRNA. In humans, this structural tRNA is *mtV*^42,43^, while *mtF* has been reported in other mammals^44^. The presence of both *mtV* and *mtF* in the monosome fraction of control MEFs suggests that both tRNAs act equally as structural components of the murine mitochondrial ribosome. In KO^MEFs^, the structural tRNA shifted towards *mtF*, probably due to the low levels of *mtV*. Furthermore, in KO^MEFs^, both structural tRNAs, but not *mtL1*, also accumulated as unprocessed transcripts in the higher molecular weight fractions (Figure 4d), suggesting they are part of a separate, larger protein-RNA complex. Thus, our results indicate that in the absence of mitoSAM, 12S and 16S rRNAs initiate early stages of assembly but stall due to a lack of processing.

### LSU assembly requires mitoSAM

The observed stalling of rRNA cleavage exposed a critical requirement for mitoSAM to facilitate entry into the ribosome assembly process. Additionally, complete monosome formation was impeded in the absence of mitochondrial methylation potential. We used cryogenic electron microscopy (cryo-EM) on mitochondrial preparations from SAMC KO^MEFs^ to visualise these assembly intermediates directly. Reconstructions from the cryo-EM dataset followed by 3D classification and refinement identified ten distinct classes representing various stages of mtLSU maturation, and no mature mitoribosomes, allowing us to establish an assembly sequence (Figure 5a and Figure S10 and S11). The multiple assembly states ranged from those with an immature central protuberance (CP) and disordered PTC to states in which 16s rRNA helix 80 (H80), 89 (H89), and 92 (H92) had acquired a near-mature conformation, despite a lack of methylation (Figure 5b). However, the PTC remained immature, preventing full mtLSU maturation and monosome formation (Figure 6).

**Figure 5.**
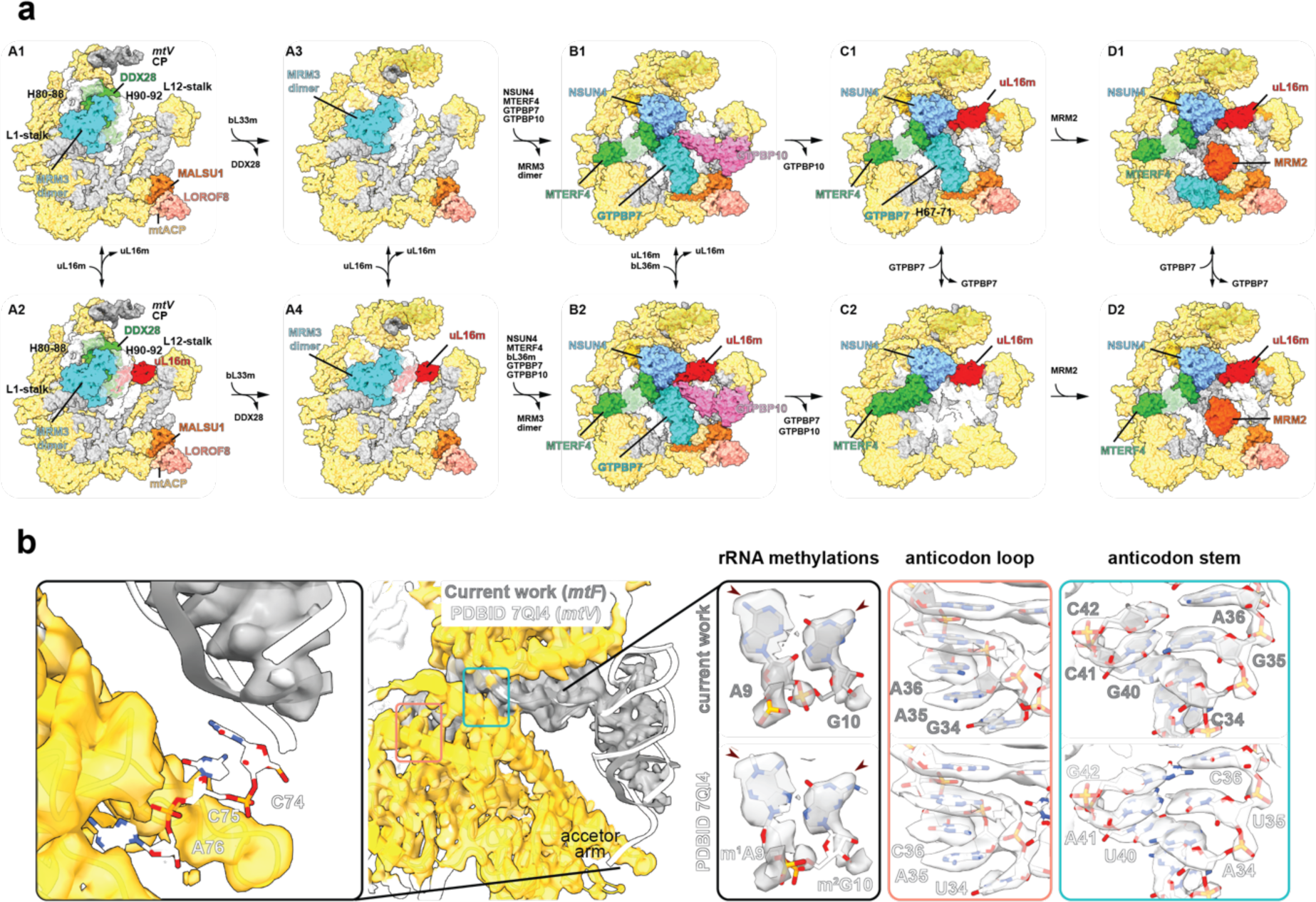
Cryo-EM structures of mouse mitochondrial LSU assembly. a, Overview of identified LSU states (A-D). 16S rRNA helices in white, CP-tRNA in grey, mature structures in yellow. Large ribosomal subunit protein uL16m (red).Biogenesis factors shown are highlighted as followed: rRNA methyltransferase 3, MRM3 dimer (cyan); DEAD box RNA helicase 28, DDX28 (green); GTP-binding protein 10, GTPBP10 (magenta); mitochondrial assembly of ribosomal large subunit protein 1, MALSU1 module (orange, salmon, pink); transcription termination factor 4, MTERF4 (dark green); 5-methylcytosine rRNA methyltransferase,NSUN4 (cyan); GTP-binding protein 7 GTPBP7 (turquoise); rRNA methyltransferase 2, MRM2 (dark orange). b, Close up of the CP showing, mtF modelled into the density map and superposed with mtV (PDB ID 7QI4 [https://doi.org/10.2210/pdb7QI4/pdb]). Zoom-in panels show a lack of 3ʹ CCA (left), lack of rRNA methylations at positions 9 and 10, comparison of anticodon loop and part of the stem of mtF and mtV against the density map.

**Figure 6.**
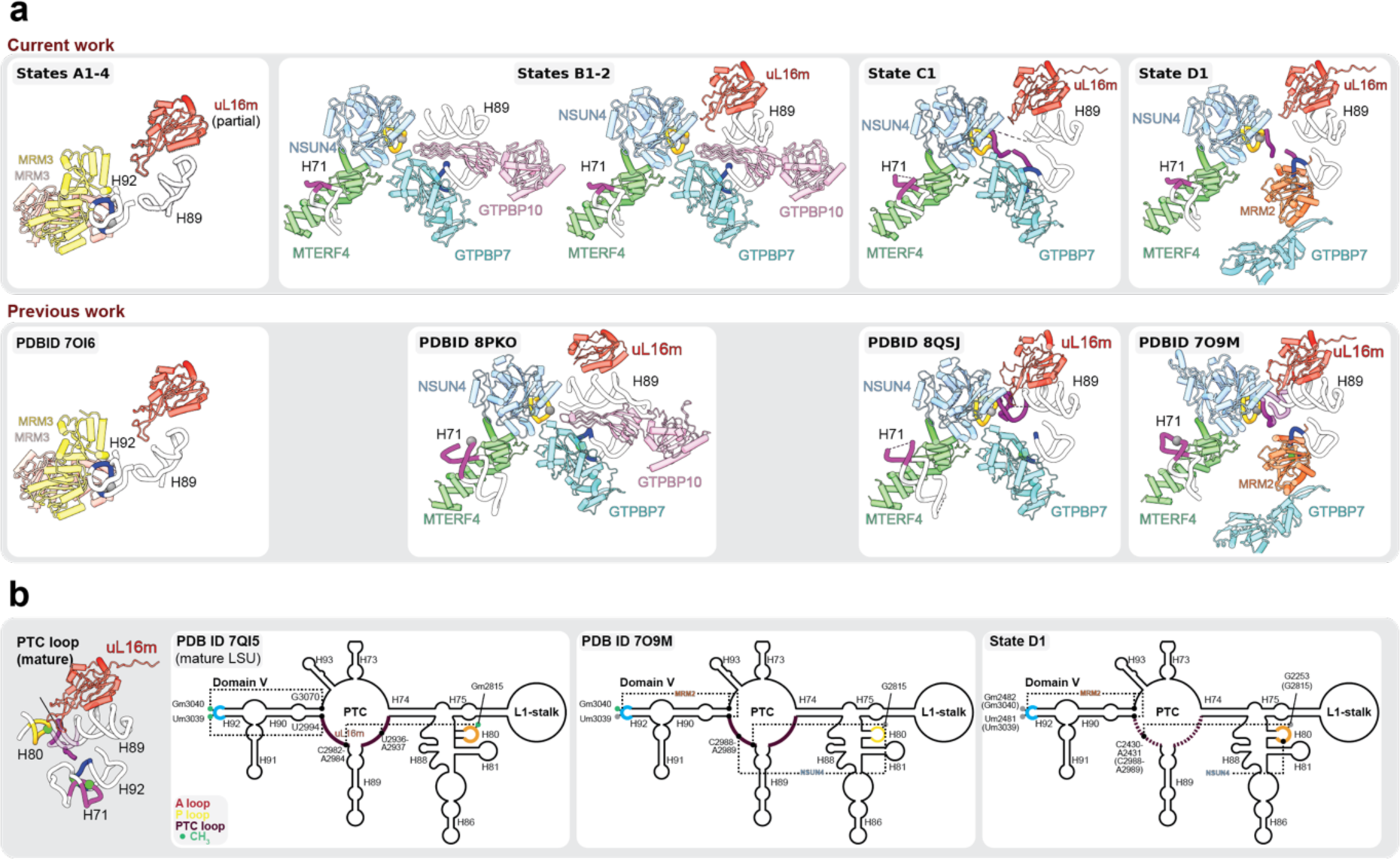
Formation of the PTC is inhibited in SAMC KO MEFs. **a**, Close-up view of PTC maturation states. Panels show LSU assembly factors bound to part of rRNA helices and loops, contributing to the PTC, namely, h67-71 (pink), h80/P loop (gold), PTC loops (purple) and h92/A loop (blue). Lower panels show comparable states from published work while upper panels depict states A1-4, states B1, B2, C1 and D1 to depict the key stages of PTC maturation observed in current work. **b**, 2D representation of domain V in mature LSU, human LSU intermediate (PDB ID 7O9M [https://doi.org/10.2210/pdb7O9M/pdb]) and state D1. Interactions made by methyl transferases MRM2 and NSUN4 are highlighted. Residues methylated in mature LSU are highlighted (methylated as light green; unmethylated as grey). Dashed lines represent predicted stabilising interactions between methylated residues. Residue numbers are shown according to mouse genome numbering and the corresponding human numbering is indicated in brackets.

The earliest states were marked by DDX28 stabilising the immature CP and an A-loop of H92 held by an MRM3 dimer, which would typically methylate G2482 (Figure 5a). Notably, despite the compromised processing of *mtF* and *mtV,* the CP contained a structural tRNA. This CP-tRNA density could be assigned as both *mtF* and *mtV*, though the mixed population demonstrated a predominance of *mtF*, supporting our ribosome gradient data (Figure 5b left panel). Surprisingly, no 3ʹCCA RNA modification was present in all mature tRNAs, suggesting that this modification is not required for incorporation into the mitochondrial ribosome.

The earliest previously reported assembly state is defined by an immature CP bound by DDX28, an MRM3 dimer, and the presence of the ObgE-like GTPase (GTPBP10) that stabilises H89 in an immature conformation^45,46^. Here, we observed four sub-states with bound MRM3 dimer, differing in (i) binding of uL16m (A2 and A4) and bL36m (A3 and A4) at the base of the L12 stalk and (ii) maturation status of the CP coupled with the dissociation of DDX28 (A3 and A4) (Figure 5a, Figure S12a). Notably, GTPBP10 did not yet appear in the assembly pathway reported here. However, in states A2 and A4, the conformation of H89 correlated with the occupancy of uL16m, where H89 is lodged against uL16m proximal to a basic patch on its surface. It is thus likely that in the absence of uL16m (states A1/A3), H89 becomes disordered or displaced (Figure S12a). Interestingly, uL16m was strongly reduced in the KO^MEFs^ mtLSU and monosome fractions in our SILAC gradient analysis, assigning uL16m a critical role during LSU assembly. States A3/A4 lacked DDX28, allowing the recruitment of bL33m and the adoption of a more mature and structured CP. The MRM3 dimer remained bound to H92 in these states, but the weak density suggested that this was only a loose interaction (Figure 5a).

In state B1 we observed (i) binding of the NSUN4-MTERF4 complex, which sequesters H68-71, (ii) recruitment of GTPBP7 and GTPBP10, (iii) the absence of uL16m, (iv) the presence of bL36m, albeit at partial occupancy, and (v) an ordered but displaced by ∼25 Å H89 to a distinct conformation stabilised by GTPBP10 (Figure 6a,b and Figure S12b). This state differed from a previous report due to the absence of uL16m (PDBID 8PKO; ref ^47^). In state B2, uL16m was stably accommodated, and H89 moved ∼14 Å to align with a basic cleft formed by uL16m and GTPBP10, acquiring its mature position. This provides insights into how these two factors promote H89 folding (Figure 6a and Figure S12b). As GTPBP10 dissociates from the maturing LSU, GTPBP7 remained bound in a closed conformation (state C1), as previously reported^48^. Finally, states D1/D2 represented the most mature states identified and characterised by the recruitment of MRM2, causing GTPBP7 to adopt an open conformation^49^.

### MitoSAM is required for PTC maturation

During translation, peptide bond formation occurs in the PTC, predominantly formed by 16S rRNA during the late stages of mtLSU assembly and involves the folding and methylation of the A and P loops^28,45,46,48–51^. The absence of mitoSAM did not prevent the recruitment of assembly factors in the ten mtLSU states reported here. Rather, we observed a partially disordered PTC. In the most mature assembly state identified here, H68-71 remained sequestered by NSUN4-MTERF4, while MRM2 was retained in its canonical position, holding the A-loop in its catalytic pocket to methylate U2481 (U3031; human mitochondrial genome numbering) (state D1 in Figure 6a). GTPBP7 was bound in a fluid conformation, and the PTC loops between H74-H89 and H89-H90 were only partly ordered (Figure 6a,b).

According to the canonical assembly pathway, state D is followed by the recruitment of GTPBP5, promoting a significant remodelling of the PTC loops and the release of MRM2 from the A-loop post-methylation^50,52,53^. The lack of GTPBP5 binding in our model can be attributed to (i) a complete absence of A-loop methylation at U2481 and G2482 (U3039 and G3040; human mitochondrial genome numbering); (ii) a clash between the unreleased MRM2 and incoming GTPBP5 due to the unmethylated A-loop; and (iii) an increased disorder of the PTC loops. Thus, in the absence of mitoSAM, multiple causes result in an unstable interface, preventing the binding of the Obg domain loops of GTPBP5.

It is not known when MRM1 methylates the P-loop during 16S rRNA maturation and LSU assembly, but it can already be observed in states containing GTPBP5 and GTPBP6^50,53^. How MRM1-dependent methylation affects PTC remodelling is also unclear, but direct contact between Gm2253 (Gm2817 in humans) and H89-H92 has been noted in human assembly intermediates (PDBID 7OF7 [https://doi.org/10.2210/pdb7of7/pdb], ref 50) and mature LSU (PDBID 7QI4 [https://doi.org/10.2210/pdb7QI4/pdb], ref 23). Our results suggest that the P-loop and the proximal PTC loop are distorted without methylation, inhibiting PTC formation.

## Discussion

Methylation is of particular importance at two sites during mitochondrial gene expression, with both rRNAs and tRNAs possessing several modifications. While several of these modifications have been linked to stabilising the decoding centre, their broader roles in function, stability, assembly, or regulation remain unclear.^23^ Here, we exclusively targeted the mitochondrial methylation potential to investigate SAM-dependent modifications within mitochondria, without affecting the responsible methyltransferases. Our findings demonstrate that methylation differentially impacts tRNA stability, with both increased and decreased steady-state levels observed.

How these modifications affect tRNA folding, and therefore stability, remains unclear. However, previous work suggests that tRNA methylation by TRMT10C is not required for the cleavage of the primary transcript by MRPP3^9,12,13^. This holds true for most gene junctions, except those involving both rRNAs, Nd2, and the mtR-Nd4/4L gene junction. The specific sensitivity of these regions is not well understood, but in the case of the rRNA gene cluster, the excessive secondary structures of the rRNAs may play a role. Therefore, structural reconstructions with these leader and trailer sequences will be essential to clarify whether these junctions fold differently. Furthermore, we identified multicistronic transcripts with tRNAs at their 5ʹ and 3ʹ termini, challenging the hierarchical cleavage of canonical gene junctions. Gene junctions not containing tRNAs were processed efficiently, consistent with previous findings that neither RNase P nor ELAC2 are responsible for their cleavage^10,54,55^.

Our work demonstrates that a decreased mitochondrial methylation potential restricts mitochondrial gene expression by impeding the processing of the ribosomal gene cluster. This, in turn, results in incomplete ribosome assembly due to the lack of *mtF* and *mtV* release from 12S and 16S rRNA. Our findings also show that the 2ʹ-*O*-methylations of 16S rRNA are crucial for the maturation of the PTC and LSU. This contrasts with recent data from human HEK 293T cells, which proposed that the 2ʹ-*O*-methylation of U2481 (U3039 in human mtDNA numbering) is dispensable for monosome formation^28^. Supporting our results, a dependency on methylation has recently been shown for the cytosolic LSU in yeast, where a single 2ʹ-*O*-methylation on the pre-60S rRNA subunit was essential for ribosome assembly^56^.

Structural analysis previously demonstrated that, besides 12S and 16S rRNA, the mitochondrial ribosome contains a third structural RNA, *mtV*^43^. Later studies reported that *mtF* substitutes *mtV* in some species and even under disease conditions^44^. Interestingly, our data suggest that in MEFs, *mtF* and *mtV* equally contribute to the murine ribosome structure. Whether this composition conveys advantages or serves regulatory purposes remains to be seen. Nevertheless, the absence of the canonical 3ʹ CCA addition of the structural CP-tRNA suggests that this modification is not required for LSU assembly and also indicates that the ribosomal gene cluster is processed and matured separately from other regions of the mitochondrial transcriptome.

We previously demonstrated that the mitochondrial SAM pool reflects its cytosolic production, providing a direct opportunity to regulate mitochondrial translation via one-carbon metabolism^35^. Here, we suggest a crucial role for mitoSAM in gating early mitochondrial ribosome biogenesis and LSU maturation. Together with the recent observation that NSUN3-mediated methylation of mtM can drive mitochondrial translation,^57^ this establishes a central role for the one-carbon cycle in controlling mitochondrial translation.

## Acknowledgements

This study was supported by grants to A.Wr. from the Swedish Research Council (VR2022-01287 and VR2023-07091), the NovoNordisk Foundation (NN0082202), the Knut and Alice Wallenberg Foundation (KAW2019.0109), the Region Stockholm (RS2022-0708), a Karolinska Institutet consolidator grant (2-190/2022), Heart and Lung Foundation (20210498) and Cancerfonden (21 1621 Pj). A.A. was supported by grants by the European Research Council (ERC-2018-StG-805230). Protein identification and quantification were carried out by the Proteomics Biomedicum Core Facility, Karolinska Institutet https://ki.se/en/mbb/proteomics-biomedicum. The authors would like to acknowledge support of the National Genomics Infrastructure (NGI) / Uppsala Genome Center and UPPMAX for providing assistance in massive parallel sequencing and computational infrastructure. Work performed at NGI / Uppsala Genome Center has been funded by RFI / VR and Science for Life Laboratory, Sweden. Cryo-EM data were collected at the Swedish National Cryo-EM Facility, SciLifeLab, Stockholm University, and at the Karolinska Institutet 3D-EM Core facility. The inhibitor of mitochondrial transcription (IMT) was a kind gift by N-G Larsson, Karolinska Institutet.

## Author Contributions

Conceptualisation, R.IC.G and A.Wr; investigation R.I.C.G; V.S., L.P-P., A.Wi., M.F.M., D.M., F.A.R., X.P., M.S., M.C., A.A., C.F., A.Wr. formal data analysis R.I.C.G., V.S., L.P-P., M.F.M., I.A., C.F., A.Wr. funding acquisition, A.We, J.R., A.A., and A.Wr; writing original draft, R.I.C.G, V.S., C.F and A.Wr; reviewing, all authors.

## Materials and correspondence

Correspondence and material requests should be addressed to Anna Wredenberg or Christoph Freyer.

## Materials and Methods

### Mouse husbandry and tissue collection

*Slc25a26* conditional KO mice (*Samc^loxp/loxp^*) were generated previously by Taconic Biosciences (Germany), by flanking exon 3 of the National Center for Biotechnology Information (NCBI) transcript NM_026255.5 with loxP sites^35^. *Samc^loxp/loxp^* mice were crossed to *Mlc1f-Cre* mice^58^ to generate skeletal muscle-specific *Slc25a26* KO (SAMC KO^SkM^) mice. All mice were maintained on a C57BL/6N genetic background and kept in individually ventilated cages at ambient room temperature of 22 – 24°C with a 12h:12h light:dark cycle and *ad libitum* access to food (Special Diet Services) and water. Groups included male and female animals unless indicated. Animal studies were approved by the local animal welfare ethics committee (Stockholm ethical committee) and performed in compliance with the national and European law.

### Generation of MEF cell lines and cell culture

MEFs were previously derived from embryonic day 13.5 embryos from intercrossed *Slc25a26*^l*oxp/lox*^ mice. In brief, isolated *Slc25a26^loxP/loxP^* MEFs were cultured at 37°C and 5% CO_2_ in medium comprising of Dulbecco’s Modified Eagle’s Medium (DMEM), high glucose, and GlutaMAX (Thermo Fisher Scientific), supplemented with 10% foetal bovine serum (ThermoFisher Scientific), and 1% penicillin/streptomycin (ThermoFisher Scientific). Immortalised *Slc25a26*^−/−^ cells were obtained by transiently expressing Cre-recombinase. Transfected cells were serially diluted to separate single cells, and subsequent clones were screened by PCR analysis. *Slc25a26^−/−^* cultured in the same medium, supplemented with uridine (50 µg/ml; Sigma-Aldrich). MEF cells were cultured in high glucose Dulbecco’s Modified Eagle Medium (DMEM) GlutaMax™, supplemented with 50 µg/mL Uridine (Sigma Aldrich), 10% Foetal Bovine Serum (ThermoFisher Scientific) and 1% Penicillin/Streptomycin (ThermoFisher Scientific). Incubator conditions were set to 37°C and 5% CO_2_.

### Mitochondrial enrichment from mouse tissue and cell lines

For enrichment of mitochondria from mouse quadricep, fresh or snap-frozen tissue was chopped until paste-like and transferred into a tissue mitochondrial isolation buffer (MIB-T) containing 225 mM Sucrose, 20 mM Tris, 1 mM EGTA. Trypsin was added at a 0.4% working concentration and samples were rotated at 4°C for 10 min. Samples were diluted with MIB-T containing 0.5% BSA and 1x Protease Inhibitor (Roche) prior to homogenisation on ice with 25 strokes on a Schuett homgen^plus^ set to 800 rpm. Tissue homogenates were differentially centrifugated at 1,000 x *g* and 11,000 x *g* and the resulting mitochondrial pellets were resuspended in MIB-T containing 0.4 mg/mL Trypsin Inhibitor before a final spin at 11,000 x *g*.

MEF cells were scraped from cell culture vessels and resuspended in ice cold Dulbecco’s Phosphate Buffered Saline then pelleted with a 5 min 800 x g spin at 4°C. The pellet was resuspended in a cell mitochondrial isolation buffer (MIB-C) containing 150 mM D-mannitol, 100 mM Tris pH7.4, 1 mM EDTA with added BSA at a working concentration of 0.1%. Cells were then homogenised on ice with 20 strokes on a Schuett homgen^plus^ set to 600 rpm. Cell homogenates were differentially centrifugated at 1,000 x *g* and 11,000 x *g* and the resulting mitochondrial pellets were resuspended in MIB-C without BSA before a final spin at 11,000 x *g*.

Mitochondrial pellets from tissue and cells were either quantified for immediate experimental use or resuspended in a freezing buffer containing 300 mM Trehalose, 10 mM Tris-HCl pH7.4, 10 mM KCl, 1mM EDTA, 0.1% BSA for long-term storage at -80°C.

### Western blot analysis

Mitochondrial pellets from MEF cells and quadricep were lysed in a mitochondrial lysis buffer (MLB) containing 50 mM KCl, 20 mM MgCl_2_, 10mM Tris pH 7.5, 1× Protease Inhibitor (Roche) and 1% Triton X-100. Mitochondrial lysates were combined with NuPAGE LDS sample buffer, loaded onto 4-12% NuPAGE Bis-Tris gels (ThermoFisher Scientific) for electrophoresis in an XCell SureLock system (ThermoFisher Scientific) and subsequently transferred to PVDF membranes using the iBlot 2 Dry Blotting System (ThermoFirsher Scientific). Membranes were blocked in 5% (w/v) milk in Phosphate Buffered Saline with 1% (v/v) Tween 20 (PBS-T; Sigma-Aldrich). Membranes we incubated in primary antibody overnight at 4°C and secondary antibody for one hour at room temperature, then developed using Clarity Western ECL substrate (Bio-Rad). List of antibodies used can be found in Table S11.

### Sucrose gradient sedimentation

Mitochondria were lysed in MLB with freshly supplemented RNase Block Ribonuclease Inhibitor (Agilent). Linear 10-30% sucrose gradients in 1x sucrose gradient buffer (50 mM KCl, 20 mM MgCl2, 10mM Tris pH 7.5, 1× Protease Inhibitor) were generated in 14 x 89 mm Ultra-Clear Centrifuge Tubes (Beckman Coulter), using a 107 Gradient Master (BioComp), followed by ultracentrifugation for 15 h at 79 000 x *g* at 4°C in an Optima L-80 XP Ultracentrifuge (Beckman Coulter SW41-Ti rotor). Starting from the top of the gradient, 25 x 450 µL fractions were collected. Fractions for western blot analysis subsequently underwent Trichloroacetic Acid (TCA) precipitation. 1/100^th^ volume of 2% Sodium Deoxycholate was added to each fraction followed by 20 minutes incubation on ice. 1/4^th^ volume of 72% TCA was added and fractions were spun at max speed in a benchtop centrifuge for 20 minutes at 4°C. Each fraction then underwent 2 washes in 100% acetone and resulting pellets were left to air dry under a fume hood. Precipitates were resuspended in NuPAGE LDS Sample Buffer freshly supplemented with 10 mM of Dithiothreitol (DTT) and either loaded directly onto an SDS-PAGE gel or frozen at -20°C for short term storage.

### In cellulo and in organello mitochondrial translation assays

MEF were seeded into 6 well plates and cultured for 48 h in high glucose Dulbecco’s Modified Eagle Medium (DMEM) GlutaMax™, supplemented with 50 µg/mL Uridine (Sigma Aldrich), 10% Foetal Bovine Serum (ThermoFisher Scientific) and 1% Penicillin/Streptomycin (ThermoFisher Scientific). Two 5 min washes with 2 mL of Cys-/Met-free medium containing DMEM High Glucose, No Glutamine, No Methionine, No Cysteine (ThermoFisher Scientific) supplemented with 10% Dialysed Foetal Bovine Serum, 1× GlutaMax, 1x Sodium Pyruvate (ThermoFisher Scientific) and 100 ug/ml Emetine. Fresh Cys-/Met-free medium was added, and the cells were incubated for a further 20 min. 200 µCi of [^35^S]-methionine/cysteine EasyTag Express protein labelling mix (Perkin-Elmer) was added to 1 mL of Cys-/Met-free medium per well and cells were incubated for 1 h at 37°C and 5% CO_2_. Following labelling, cells were washed three times with PBS, then harvested using cell scrapers and pelleted with a 5 min, 5,000 x *g* spin at 4°C. Cell pellets were resuspended in PBS supplemented with 1x Protease Inhibitor (Roche) and 50U of Benzonase (Sigma Aldrich) and then frozen overnight at -20°C to enable freeze-thaw lysis. Protein content of lysates was determined with a Pierce BCA assay (ThermoFisher Scientific), according to the manufacturers protocol, and mixed with NuPAGE LDS sample buffer, ready for SDS-PAGE.

Mitochondria from tissue quadricep were enriched through differential centrifugation in MIB-T. 500 ug of mitochondria were resuspended in 750 µg of translation Buffer (100 mM Mannitol, 10 mM Sodium Succinate Dibasic Hexahydrate, 80 mM Potassium Chloride, 5 mM Sodium Chloride, 1mM Potassium Phosphate Dibasic, 25 mM Hepes, 60 µg/mL 17 Amino Acid Mix (Ala, Arg, Asp, Asn, Glu Gln, Gly, His, Ile, Leu, Lys, Phe, Pro, Ser, Thr, Trp, Val), 60 µg/mL Cysteine, 60 µg/mL Tyrosine, 5 mM ATP, 200 µM GTP, 6 mM Creatine Phosphate, 60 µg/mL Creatine Kinase and 200 µg/mL Emetine) with 150 µCi of EasyTag L-[^35^S]-Methionine. Samples were incubated for a one-hour pulse labelling, then chase samples were washed with isotope-free Translation Buffer, supplemented with 60 µg/mL cold methionine, and incubated for a further 3 h. Samples were washed one final time, prior to resuspension in NuPAGE LDS sample buffer.

Samples from *in cellulo* or *in organello* labelling were loaded onto a 12% NuPAGE Bis-Tris gel (ThermoFisher Scientific) and electrophoresed at 150V in an XCell SureLock system (ThermoFisher Scientific) for approximately 1 h. The gel was incubated in Imperial Protein Stain (ThermoFisher Scientific) and imaged to assess loading, prior to fixing in 20% Methanol, 7% Acetic acid, 3% Glycerol for 1 h at room temperature. After fixation the gel was dried under vacuum at 60°C for 2 h, then placed to expose in a Fuji Film Phosphor Screen cassette. The resulting signal was detected using the Typhoon FLA 7000 Phosphorimager (GE Healthcare).

### RNA isolation, reverse transcription, and quantitative PCR

RNA was isolated using RNeasy Fibrous Tissue Mini Kit (Qiagen), RNeasy Mini Kit (Qiagen), or TRIzol reagent (ThermoFisher Scientific). RNA isolated for downstream qRT-PCR or sequencing experiments were subjected to DNase treatment either on-column (Quiagen RNase Free DNase Set) or column-free (ThermoFisher Scientific TURBO DNA free kit). RNA was reverse-transcribed with the High-Capacity cDNA Reverse Transcription Kit (Applied Biosystems). qRT-PCR was performed using TaqMan probes and TaqMan Universal Master Mix II (ThermoFisher Scientific) on a QuantStudio 6 System. B-actin was used as a loading control for normalisation of target genes.

### Bisulphite pyrosequencing

One µg of total RNA per sample was bisulfite converted with the EZ RNA methylation kit, following manufacturer’s instructions (Zymo Research). Reverse transcription and PCR amplification was performed with a PyroMark RT kit (QIAGEN) using primers optimised for the bisulfite modifications with one primer biotinylated. cDNA amplicons were enriched with streptavidin-coupled sepharose beads before following the manufacturer’s recommendations for pyrosequencing.

### Northern blot analysis

An appropriate amount (0.5 – 2 µg) of total RNA was separated in 1% MOPS-formaldehyde agarose gels and transferred to Hybond-N + membranes (GE Healthcare). Membranes were exposed to either randomly [^32^P]-labelled dsDNA probes, [^32^P]-labelled strand-specific RNA probes or with [^32^P]-end labelled oligonucleotide probes, using RapidHyb (Sigma Aldrich). Membranes were exposed to a PhosphorImager screen, and the signal was quantified using a Typhoon FLA7000 system and ImageQuant TL 8.1 software (GE Healthcare). Primers used to generate dsDNA and oligonucleotide probes are listed in Table S11.

### Sample preparation for Nanopore direct long-read RNA sequencing

RNA was isolated from isolated mitochondria from MEFs or skeletal muscle using the RNeasy Mini Kit (Qiagen) or RNeasy Fibrous Tissue Mini Kit (Qiagen) kits, following the manufacturer’s instructions. Input quality control (QC) of samples was performed on an Agilent 2100 Bioanalyzer, using the Eukaryote Total RNA Nano kit to evaluate RIN values and concentration (Agilent). Dynabead polyA enrichment was performed on 6-18ug of whole RNA, using the Invitrogen Dynabeads™ mRNA Purification Kit (ThermoFisherScientific Cat No 61006). RNA libraries were prepared as described in the Oxford Nanopore “Direct RNA sequencing (SQK-RNA002)” protocol, version DRS_9080_v2_revU_14Aug2019, using the Oxford Nanopore Direct RNA Sequencing Kit (SQK-RNA002). 125-439 ng of polyA enriched RNA was used for reverse transcription, followed by ligation of sequencing adapters. QC of the libraries was performed with the Qubit dsDNA HS kit (ThermoFisher Scientific). Finally, the samples were loaded on SpotON flow cells (FLO-MIN106), using the Flow Cell Priming Kit (EXP-FLP002), and sequenced on the Oxford Nanopore MinION system.

### Nanopore data analysis

Fast5 files were merged using the multi_to_single_fast5 function from the ont_fast5_api toolkit (https://github.com/nanoporetech/ont_fast5_api/tree/master). The merged files were then re-basecalled using Guppy v4.4.1 with the following parameters: --flowcell FLO-MIN106, --kit SQK-RNA002, --recursive, --fast5_out, and --qscore_filtering 7. The resulting FASTQ files were mapped to the mitochondrial genome in the GRCm38 reference using Minimap2 v2.17 with parameters: -ax splice -uf -k14^59^. The SAM files generated from this mapping were converted to BAM format, then sorted and indexed using Samtools v1.10^60^. Next, the BAM files were converted to BED format using the bamtobed function from Bedtools v2.29.2^61^. The resulting BED files with same age and genotype were merged, processed, and visualised in R v4.3.1 using the ggplot2 package v3.4.3 for visualisation.

### Stable isotope labelling

KO and Control MEFs were grown for 19 days in DMEM for SILAC (ThermoFisher Scientific) supplemented with 10% Dialysed FBS, 200 mg/mL Proline (Sigma Aldrich), 50 µ)g/mL Uridine (Sigma Aldrich) and either heavy (^13^C_6_, ^15^N_4_) or light L-Arginine and heavy (^13^C_6_,^15^N_2_) or light L-Lysine, followed by mitochondrial enrichment.

### Sucrose gradient fraction proteomics

Mitochondrial pellets from labelled KO and Control MEF cells were enriched in MIB-C and quantified, then 1.5 mg of heavy Arg/Lys KO and light Arg/Lys Control or heavy Arg/Lys Control and light Arg/Lys KO were combined, to give two replicate experiments each with a total of 3 mg of mitochondrial content. Merged mitochondria were lysed in MLB freshly supplemented with RNase Block (Agilent), as previously described for western blot analysis The mixed mitochondrial lysate was loaded onto a linear 10-30% sucrose gradient and subjected to ultracentrifugation sedimentation. Fractions of 450 µL were taken and ¼ of fractions 1-19 were TCA precipitated for SDS-PAGE and Western blotting to establish the mitoribosome sedimentation pattern. The remaining volume of fractions 9-11, 12-14 and 15-17, corresponding to mtSSU, mtLSU and monosome, were pooled and precipitated with three volumes of Ethanol. Protein pellets were resuspended in 20 µL of 6 M guanidine hydrochloride (GdmCL), 10 mM Tris-(2-carboxyethyl)phosphine hydrochloride (TCEP), 40 mM 2-chloroaceteamide (CAA), 100 mM Tris– HCl, pH 8.5. Overnight digestion and peptide cleaning was performed as described previously^62^. One third of the sample was used for LC-MS/MS analysis.

### LC-MS/MS analysis

Peptides were separated on a 40 cm, 75 μm internal diameter packed emitter column (Coann emitter from MS Wil, Poroshell EC C18 2.7 micron medium from Agilent) using an EASY-nLC 1200 (ThermoFisher Scientific). The column was maintained at 50°C. Buffer A and B were 0.1% formic acid in water and 0.1% formic acid in 80% acetonitrile, respectively. Peptides were separated at a flow rate of 300 nl / min, on a gradient from 6% to 31% buffer B for 57 min, from 31% to 44% buffer B for 5 min, followed by a higher organic wash. Eluting peptides were analysed on a Orbitrap Fusion Tribrid mass spectrometer (ThermoFisher Scientific). Peptide precursor m/z measurements were carried out at 60000 resolution in the 350 to 1500 m/z range. The most intense precursors with charge state from 2 to 7 only were selected for HCD fragmentation using an isolation window of 1.6 and 27% normalized collision energy. The cycle time was set to 1 sec. The m/z values of the peptide fragments were measured at a resolution of 30000 using an AGC target of 2e5 and 54 ms maximum injection time. Upon fragmentation, precursors were put on a dynamic exclusion list for 45 sec.

### Protein identification and quantification

The raw data were analysed with MaxQuant version 1.6.1.0^63^. Peptide fragmentation spectra were searched against the canonical and sequences of the *Mus musculus* reference proteome (proteome ID UP000000589, downloaded December 2018 from UniProt). Methionine oxidation and protein N-terminal acetylation were set as variable modifications; cysteine carbamidomethylation was set as fixed modification. Multiplicity was set to two; Arg10 and Lys8 were set as heavy labels. The digestion parameters were set to “specific” and “Trypsin/P,” The minimum number of peptides and razor peptides for protein identification was 1; the minimum number of unique peptides was 0. Protein identification was performed at a peptide spectrum matches and protein false discovery rate of 0.01. The “second peptide” option was on. Differential abundance analysis was performed using limma, version 3.34.9^64^ in R, version 3.4.3^65^. Protein groups annotated as Potential contaminant, Reverse, or Only identified by site were removed prior to the statistical analysis. The mass spectrometry proteomics data have been deposited to the ProteomeXchange Consortium via the PRIDE partner repository^66^.

### Cryo-EM sample preparation

Mitochondrial pellets from MEF cells were freshly enriched then further clarified using a step gradient of 1M and 1.5M Sucrose in a buffer of 20 mM Tris and 1mM EDTA. The bi-layer step gradients were prepared in 14 x 89 mm Ultra-Clear Centrifuge Tubes (Beckman Coulter). Mitochondria were resuspended in MIB-C buffer and loaded onto the top of the step gradient, then spun in an Optima L-80 XP Ultracentrifuge (Beckman Coulter SW41-Ti rotor). at 25,000 rpm for 1 h at 4°C. Clarified mitochondria sedimented as a band at the interface of 1M and 1.5M sucrose, which was then collected and diluted with an equal volume of 10 mM Tris-HCl pH 7.4. The samples were subjected to a final 10 min cold spin at 11,000 x *g* and then resuspended in a freezing buffer containing 300 mM Trehalose, 10 mM Tris-HCl pH 7.4, 10 mM KCl, 1mM EDTA, 0.1% BSA for long-term storage at -80°C.

Mitochondria were thawed and resuspended in 2% β-DDM (*n*-dodecyl β-d-maltoside), 25 mM HEPES, 10 mM Mg(OAc)_2_, 50 mM KCl, 1 mM DTT and protease inhibitor cocktail (Roche) incubated on ice. The mitochondrial suspension was then subjected to homogenisation with 10-20 manually administered strokes in a glass homogeniser. The solution was kept on a rocker at 4°C for 20 min. The lysate was layered on top of 0.8 M sucrose solution buffered with 25 mM HEPES, 10 mM Mg(OAc)_2_, 1 mM DTT and 1% β-DDM in a TLA120.2 thick-walled tube and subjected to ultracentrifugation at 100 000 rpm in a TLA 120.2 rotor (Beckmann Coulter) for 1 hour. The resulting pellet was then gently washed and resuspended in 25 mM HEPES, 10 mM Mg(OAc)_2_, 50 mM KCl, 1 mM DTT and 0.02 % β-DDM. The solution was clarified twice by centrifugation at 15000 x *g* for 15 min and the final solution was used for cryo-EM grid preparation.

To capture more assembly states including those with bound GTPases, mitochondria were lysed using the same protocol as above with the addition RNase inhibitor and 0.5 mM GMPPNP (a nonhydrolysable GTP analog to the lysis and resuspension buffers. Further, to prevent potential degradation of sensitive complexes due to blotting and vitrification, 0.5 mM BS3 (a crosslinking agent) was added 10 min prior to grid freezing. The sample was then clarified with centrifugation at 15 000 x *g* for 2 min before being used. A volume of 3 µL sample (A_260_ 1.64) was placed on a glow discharged (for 30 sec at 20 mA) Quantifoil Au 300 mesh R2/1 grid coated with ∼3 nm thick continuous carbon support in a controlled environment of 100 % humidity and 4°C temperature using Vitrobot mKIV (FEI/Thermo Fisher). After 30 sec of incubation, the excess sample was blotted off for 3 sec and subjected to vitrification in liquid ethane. The frozen grids were then stored in liquid nitrogen for data collection.

### Cryo-EM data acquisition and image processing

Data acquisition was carried out using a FEI Titan Krios (FEI/ThermoFisher Scientific) 300 kV electron microscope equipped with a Gatan K3 detector. For the untreated sample, a total of 65 125 movies (dataset 1) were collected (35 frames each) at a defocus range of -0.5 to -1.8, an electron dose rate of 1.05 e/frame/Å^2^ at a pixel size of 0.846 Å (105 000x magnification). For the treated sample, two cryo-EM datasets (2 and 3) were acquired. Dataset 2 had a total of 47 960 movies (35 frames each) collected at a defocus range of -0.5 to -1.8, electron dose rate of 1.05 e/frame/ Å^2^ with pixel size of 0.846 Å^2^. Dataset 3 had a total of 60 000 movies (35 frames each) collected at a defocus range of -0.5 to -1.8, electron dose rate of 1 e/frame/ Å^2^ with pixel size of 0.825 Å.

Motion correction was carried out in Relion 3.1.1^67^ using its implementation of MotionCorr 2.0^68^. The motion corrected micrographs were used for CTF estimation with GCTF^69^. Particles were exhaustively picked by blob picking and separately with SSU, LSU and monosome class averages used as references. The picked particles were extracted at a box size of 540 pixels binned to 180 pixels and subjected to 2D classification. The particle images from classes corresponding to the large subunit/assembly states were pooled to produce a consensus map with 3D-autorefinement. The data was further cleaned up with 3D classification with local angular search range of +/-1.8° and the junk images were rejected. These were then subjected to per-particle CTF correction (beam-tilt, per-particle defocus, per-micrograph astigmatism) followed by Bayesian Polishing and a second round of CTF refinement (beam-tilt, per-particle defocus, per-micrograph astigmatism, trefoil and fourth-order aberrations, magnification anisotropy). At this stage, the particles from datasets 2 and 3 were pooled together. The particle images were now binned again 180 pixels and subjected to a series of unaligned focussed 3D classifications with signal subtraction (FCwSS) on the interface and around factor binding regions to isolate distinct mtLSU assembly intermediates. Particles from dataset 1 (native) classified into MRM3-dimer states (unsorted states A1 to A4), NSUN4-MTERF4 bound states with/without MRM2 and GTPBP7 in open conformation (states C2, D1+D2). Particles from datasets 2 and 3 classified into two states with GTPBP7 (closed conformation) (state C1) and GTPBP7 (closed conformation) and GTPBP10 (unsorted states B1 and B2). The particles corresponding to identified assembly intermediates were then unbinned to 540 pixel box size and used for 3D auto-refinement to produce the final assembly state maps (Figure S12). MRM3-dimer bound states were further sorted into four subsets based on the occupancy of DDX28, followed by uL16m by performing FCwSS on the central protuberance and uL16m, respectively (sorted states A1, A2, A3 and A4). Similarly, unaligned FCwSS was done by masking uL16m to sort GTPBP10 bound states based on the occupancy of uL16m (sorted states B1 and B2). MRM2 bound state (D1+D2) was classified further into GTPBP7 bound (state D1) and unbound states (state D2) by performing masked FCwSS on GTPBP7 binding region. The maps were sharpened with an appropriate *B*-factor and local-resolution filtered (Figure S12). The maps with MRM3 dimer were at relatively lower resolution and suffered from substantial anisotropy. The corresponding particle images were exported to CryoSparc v4.1^70^ and subjected to nonuniform-refinement which significantly reduced map anisotropy. The maps thus obtained were then used for model building. For generating a good quality map for CP-tRNA, particle images from NSUN4-MTERF4-bound states (in which the CP has matured) were pooled and subjected to 3D autorefinement yielding a map of ∼3.1 Å overall resolution. The map was then *B*-factor sharpened and local-resolution filtered to be used for the identification and modelling of the CP-tRNA species. The data acquisition parameters and data processing details are listed in Table S12.

### Model building and refinement

Manual model building was done using PDB 7O9M^71^ [https://doi.org/10.2210/pdb7O9M/pdb] as the starting template for NSUN4-MTERF4 bound states and PDB 7OI6^72^ [https://doi.org/10.2210/pdb7OI6/pdb] as the template for MRM3 dimer bound states. First, the models were rigid-body fitted into the respective maps in UCSF Chimera v1.14^73^. Next, each chain was individually rigid-body fitted into the individual maps in coot v0.9.1^74^. The sequence for mouse 16S rRNA and mitoribosomal proteins were taken from NC_005089. The sequence of the rRNA and protein chains in the structures were mutated in coot v0.9.1 using the inbuilt ’Align and mutate’ function. The alignment and mutations generated were manually checked against the map and corrected wherever required.

CP-tRNA model was built by docking the model of aminoacylated *mtV* (from PDB 7QI4^75^ [https://doi.org/10.2210/pdb7QI4/pdb] into the local resolution filtered map of the central protuberance. The sequence was then mutated to mouse *mtV* and real-space refined into the map with significant remodelling of the elbow region. The density revealed an incomplete match between the bases and the density. Mutating the sequence to *mtF* provided a much better agreement of the bases with the density (Figure 5b). However, there is an extra density at the acceptor arm of the tRNA which can accommodate a single nucleotide. Since, *mtV* is one residue longer than *mtF*, it potentially indicates the presence of *mtV* at the CP, however at a lower occupancy as compared to *mtF.* Therefore, a molecule of *mtF* was modelled here and retained in all assembly states in this work. There was no density observed for 3’ CCA (Figure 5b).

For MRM3-bound states, the chains for DDX28, uL16m and bL36m were removed in the states which lacked the corresponding density. None of these states had a density for GTPBP10 contrary to the previous observation. Hence the chain was removed and H89 was remodelled depending on the presence or absence of uL16m. Some densities were left unmodelled as the poor resolution did not allow identification. The chains were then subjected to self-restrained real-space refinement individually. For NSUN4-MTERF4-bound states (B1-D2), the models were subjected to real-space refinement and agreement with the map was checked and ensured in a residue-by-residue manner for most of the chains with sufficient local resolution. Specifically, for states B1 and B2, the initial model for GTPBP10 was taken from AlphaFold Protein Structure Database ^76,77^ (AF-Q8K013-F1) and fitted into the map. Due to relatively poor local resolution the model was fitted with coot generated reference-restrained real-space refinement to ensure a good agreement with the density on the level of secondary structure. Model for GTPBP7 was taken from PDB 7PD3^48^ [https://doi.org/10.2210/pdb7PD3/pdb]. uL16m was removed and H89 was remodelled to agree with the density maps of states B1 and B2. A molecule of GMPPNP was placed in the binding pockets of GTPBP7 and GTPBP10. For state C1, GTPBP10 was removed. For state D1, particles from states D1 and D2 were pooled to improve the quality of the map; GTPBP7 was partly retained in the open conformation as in PDB 7O9M [https://doi.org/10.2210/pdb7O9M/pdb]. A molecule of SAH was placed in the binding pocket MRM2 as a distinct density could be observed. Real-space refinement of the models was carried out with torsion and Ramachandran restraints. Restraints for ligands and rRNA modifications were generated from CCP4 7.0 library^78^ and imported into coot for model building and refinement. Models were not built for states A4 and C2 due to low resolution or high anisotropy.

The models were then hydrogenated using ReadySet from the PHENIX suite^79^. The output models were further refined against the respective *B*-factor sharpened, local-resolution filtered maps by carrying out energy minimization and ADP (atomic displacement parameter) estimation with rotamer and Ramachandran restraints using Phenix.real_space_refine v1.21^79^ followed by validation with MolProbity^80^.

