## Extended data Figures 1-12 for "The mitochondrial methylation potential gates mitoribosome assembly"

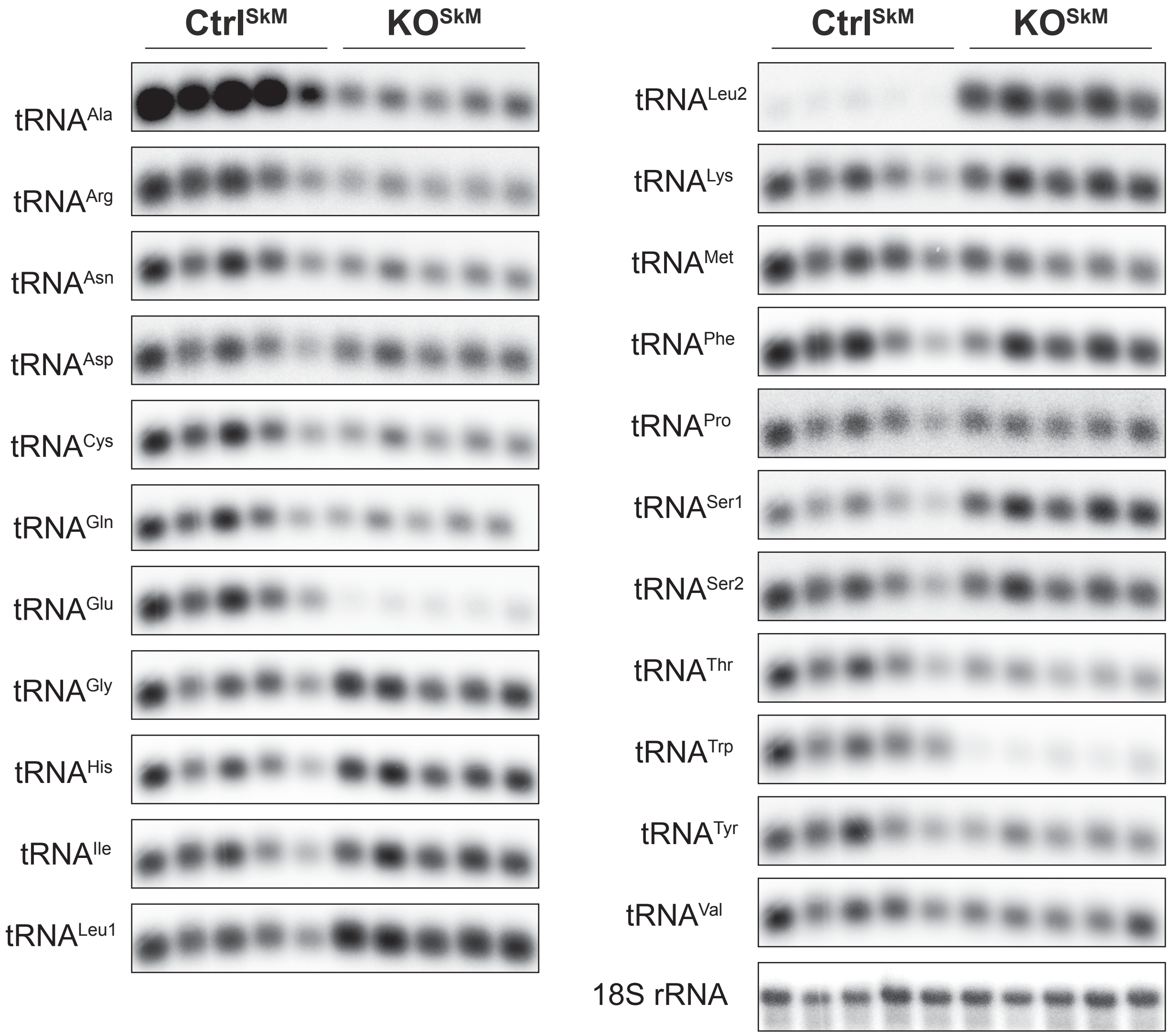


**Supplementary Fig. 1. Transcript levels**

mt-tRNA steady-state levels determiend by Northern blot analysis in quadriceps from control ($Ctrl^{SkM}$) and SAMC KO ($KO^{SkM}$) mice at 12 weeks of age. Oligo probes were used against transcripts as indicated. 18S rRNA was used as loading control.

**a**

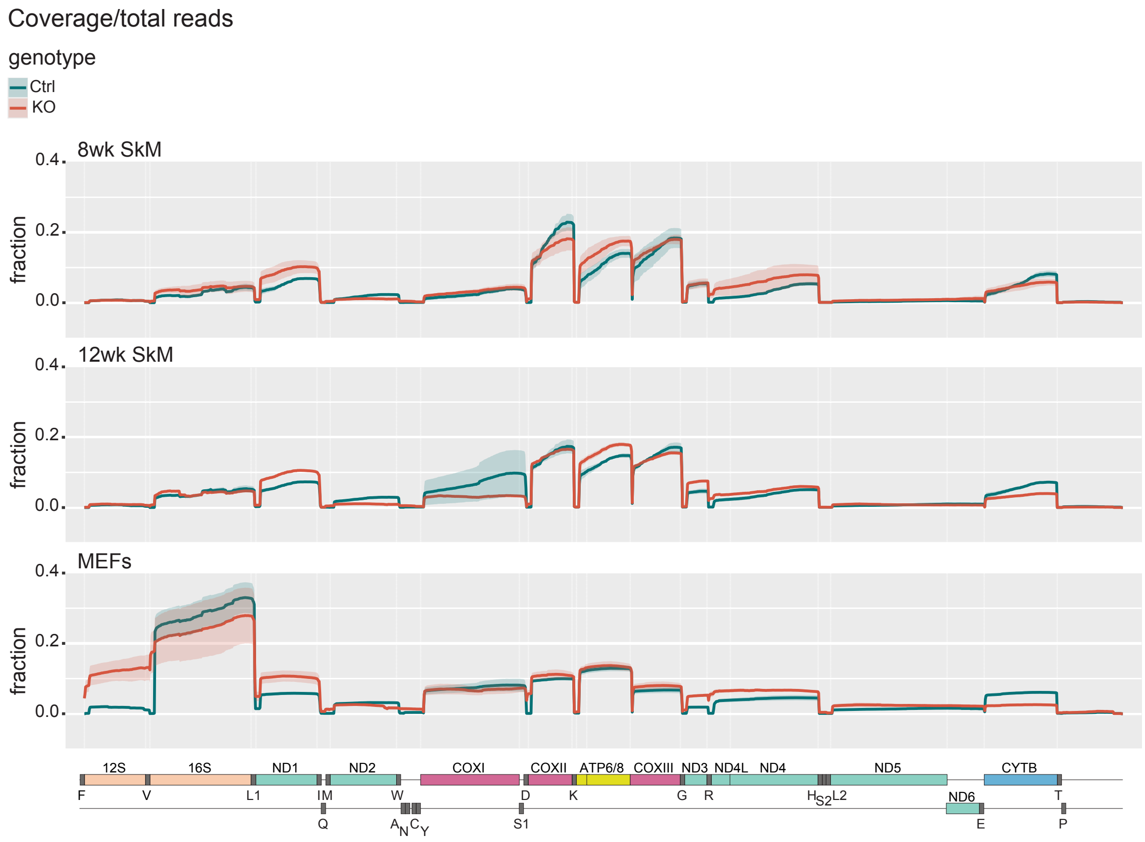


**b**

| | 8wk | | 12wk | | | |
|---|---|---|---|---|---|---|
| | Ctrl$^{SkM}$ | KO$^{SkM}$ | Ctrl$^{SkM}$ | KO$^{SkM}$ | Ctrl$^{MEF}$ | KO$^{MEF}$ |
| 12S | 98,3 | 85.6 | 98,7 | 63,5 | 91,4 | 5,8 |
| 16S | 49,9 | 43,5 | 41,9 | 42,5 | 14,9 | 5,9 |
| Nd1 | 79,1 | 74,7 | 74,4 | 68,5 | 52,2 | 40,0 |

### Supplementary Fig. 2.

**a**, Coverage plots for mtDNA from ONT sequencing of skeletal muscle samples at 8 and 12 weeks of age and MEFs. Coverage caluclated as a fraction of total reads. KO samples plotted in red, Control samples plotted in green. **b**, Percentage of completely processed (at both 5’ and 3’ ends) 12S (RNR1), 16S (RNR2) and ND1 trancripts within each sample group.

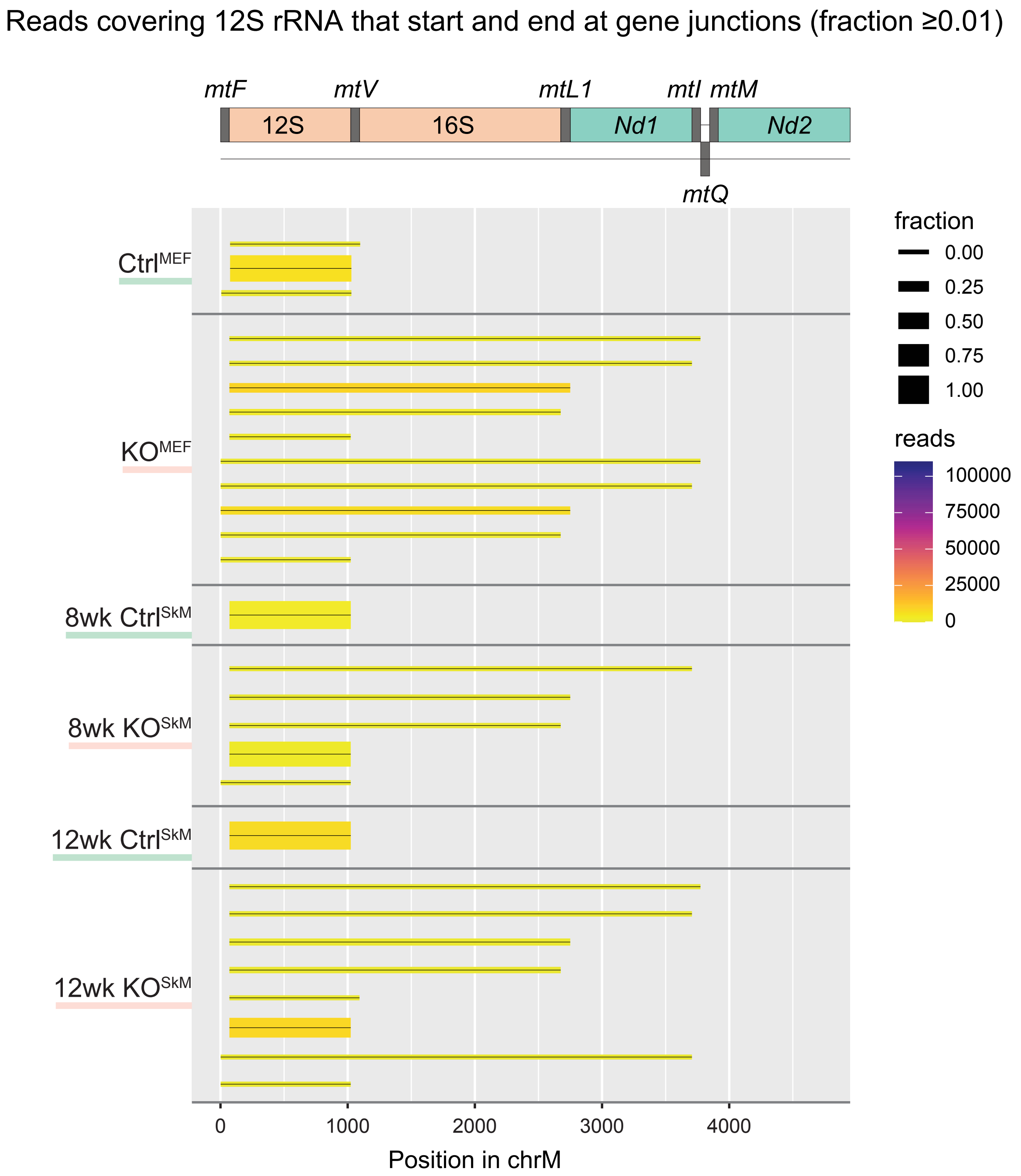


**Supplementary Fig. 3**
Start and end sites of transcripts containing 12S rRNA and a ≥0.01 fraction of total reads.

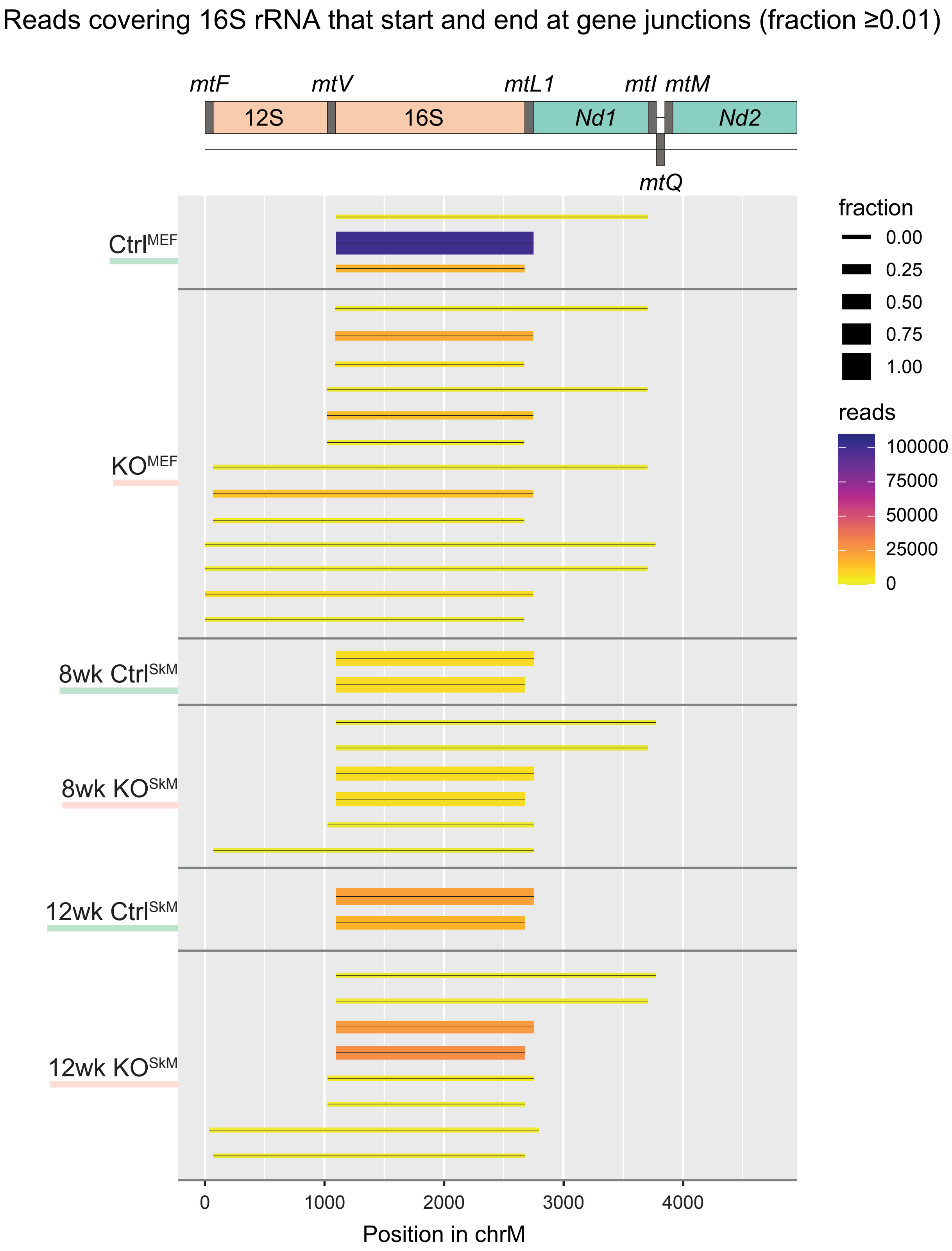


**Supplementary Fig. 4**
Start and end sites of transcripts containing 16S rRNA and a ≥0.01 fraction of total reads.

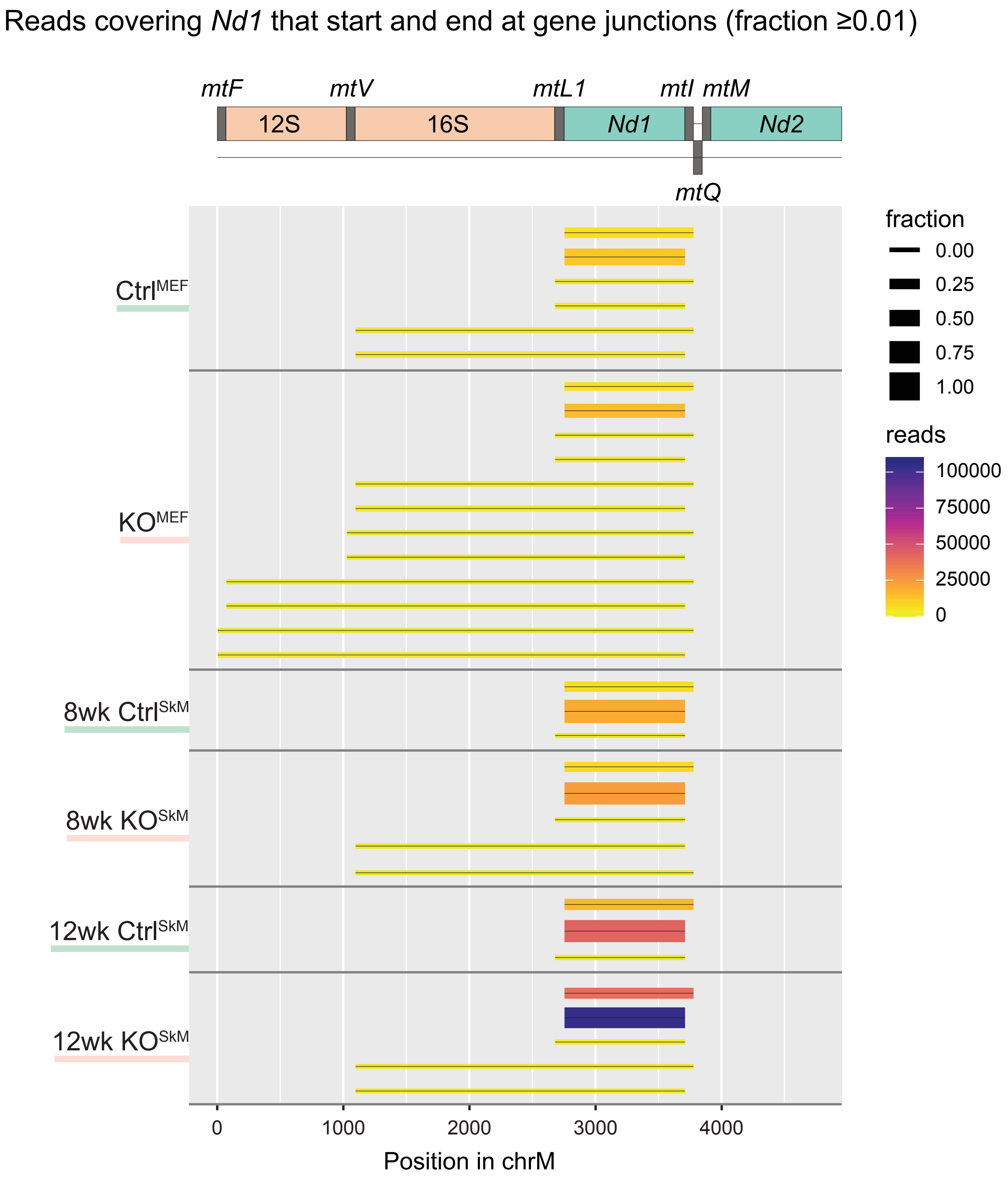


**Supplementary Fig. 5**
Start and end sites of transcripts containing *Nd1* and a ≥0.01 fraction of total reads.

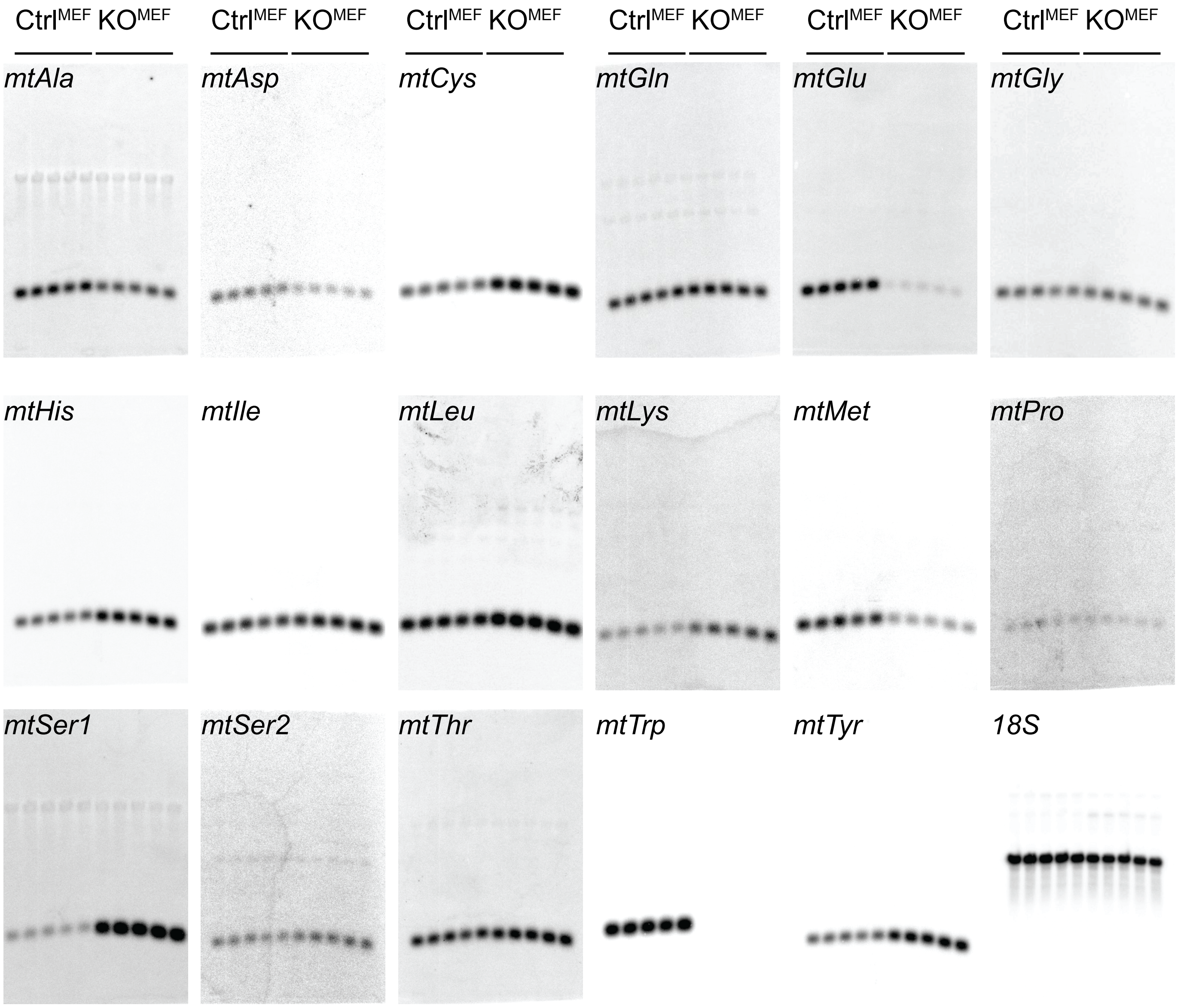


**Supplementary Fig. 6**

Northern blot analysis of Ctrl and SAM KO MEFs, using probes against tRNAs as indicated. 18S is used as loading control.

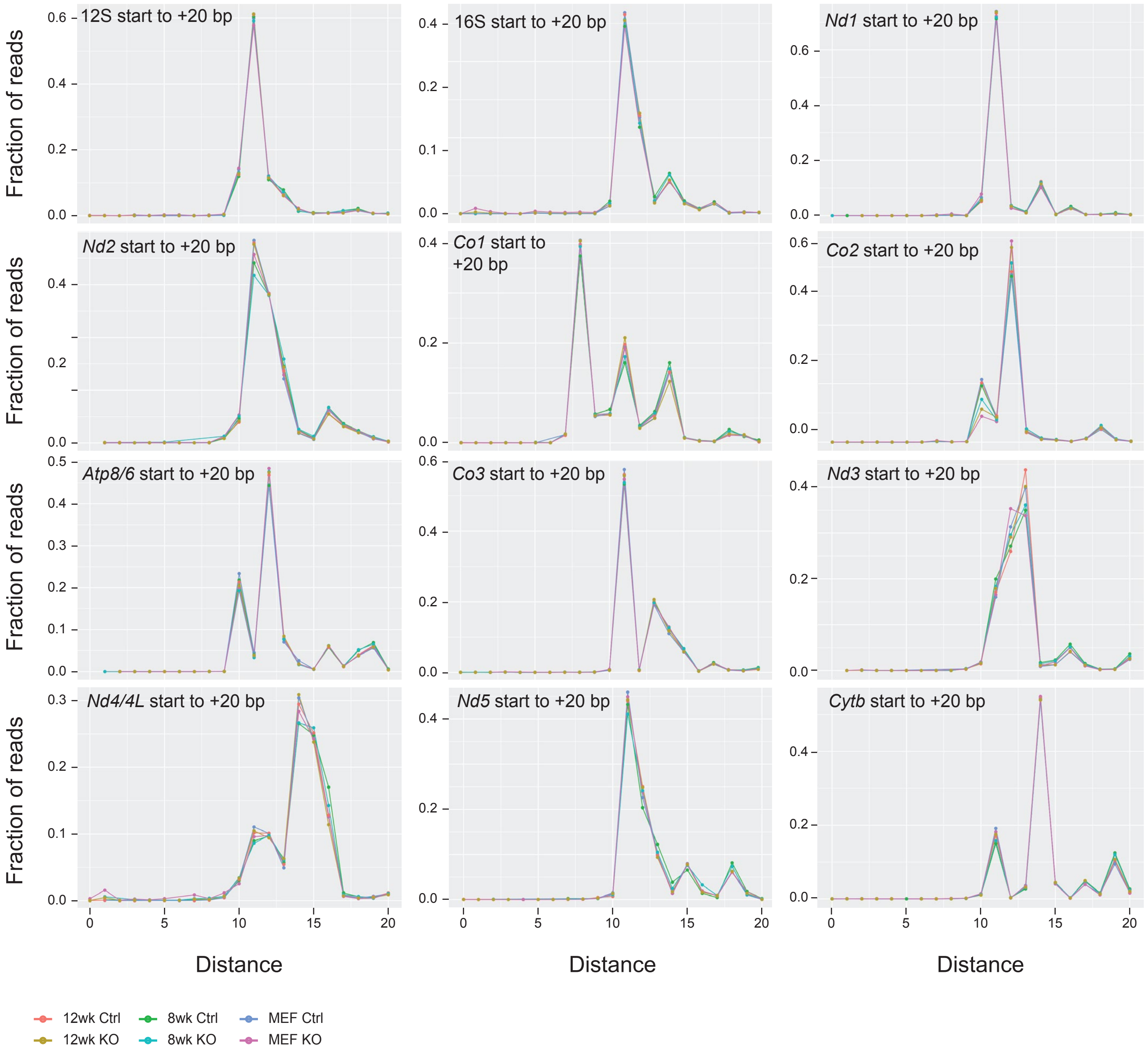


### Supplementary Fig. 7

Start site frequency of mature transcripts in ONT data set as indicated. Distance from the anotated start site is shown.

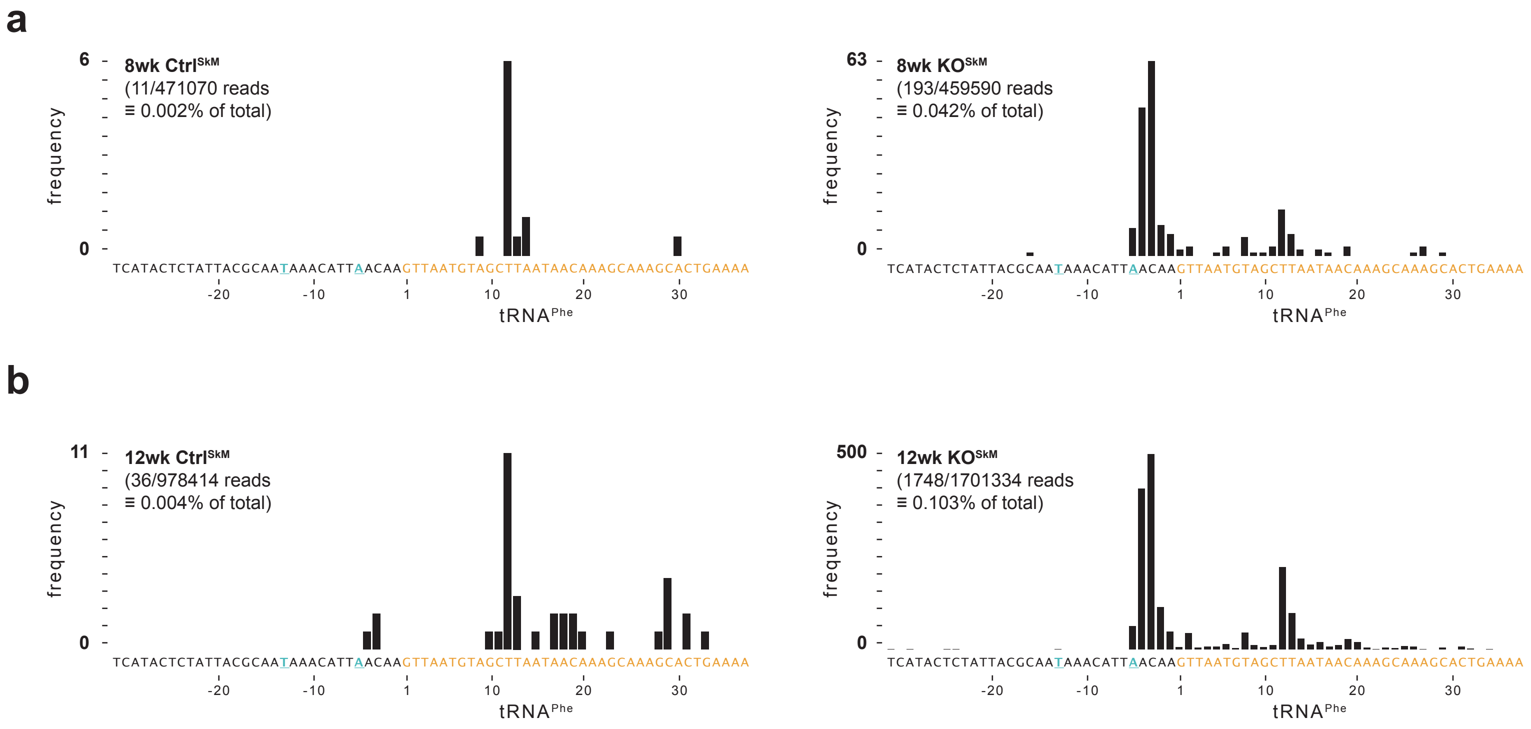


### Supplementary Fig. 8

Number of start sites for reads containing mt-tRNA$^{Phe}$ as detected by ONT analysis in quadriceps preparations from **a**, 8-week-old and **b**, 12-week-old animals.

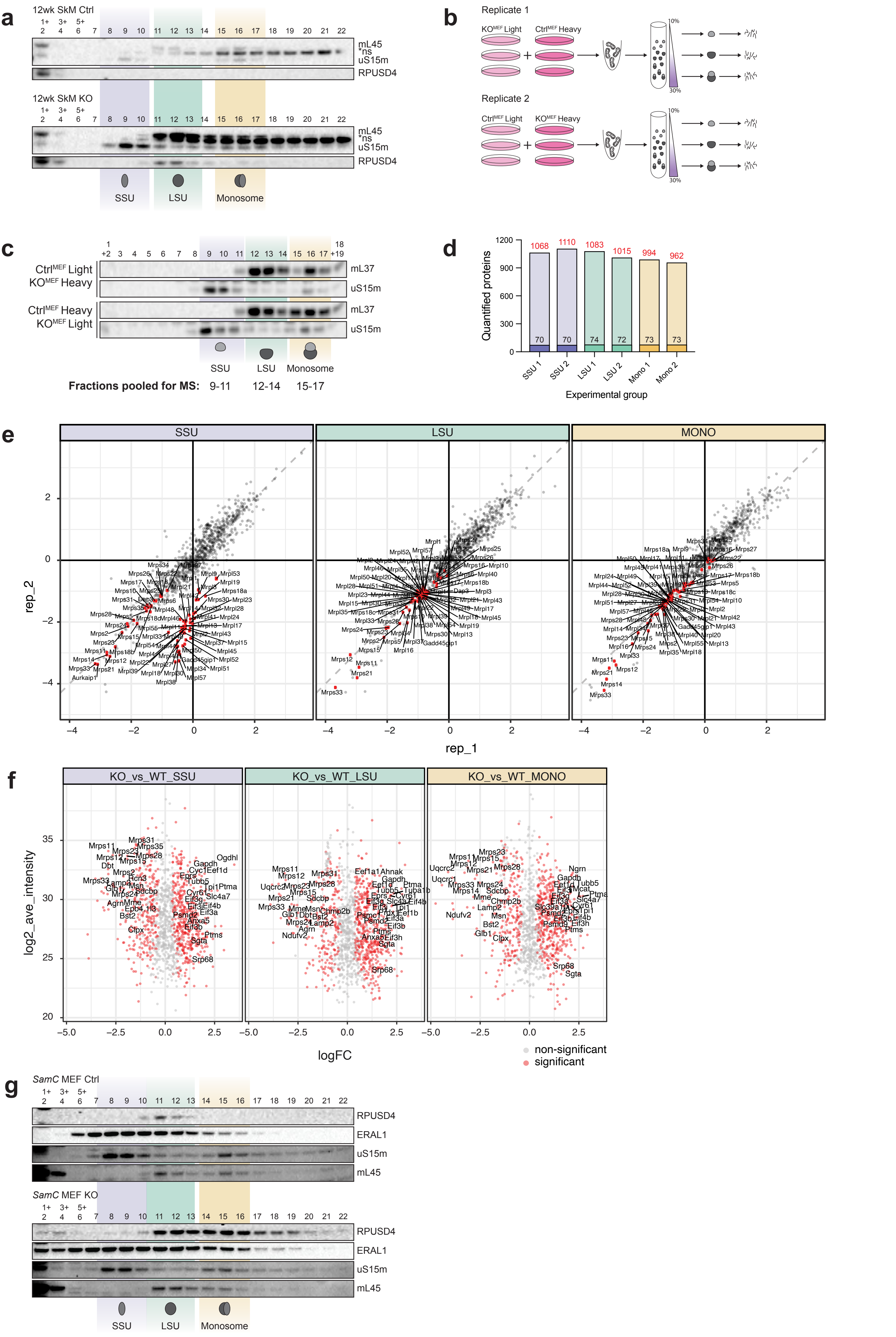


### Supplementary Fig. 9

**a**, Western blot analysis of sucrose gradient fractions from 12-week-old quadriceps mitochondrial preparations, decorated with antibodies as indicated. *ns = non-specific band. **b**, Scheme of SILAC labelling. **c**, Western blot analysis of pooled samples after SILAC labelling and sucrose gradient separation decorated with antibodies as indicated. **d**, Number of peptides identifed per sample. Total number is shown in red, mitoribosome-specific numbers (dark) are shown in white. **e**, Ratios of SILAC proteomes (Ctrl vs SAM$^{KO}$) in three different fractions as indicated. **f**, Differential expression of SILAC-labelled peptides in the indicated genotypes represented as MA plots. **g**, Western blot analysis of sucrose gradient fractions from MEF mitochondrial preparations, decorated with antibodies as indicated.

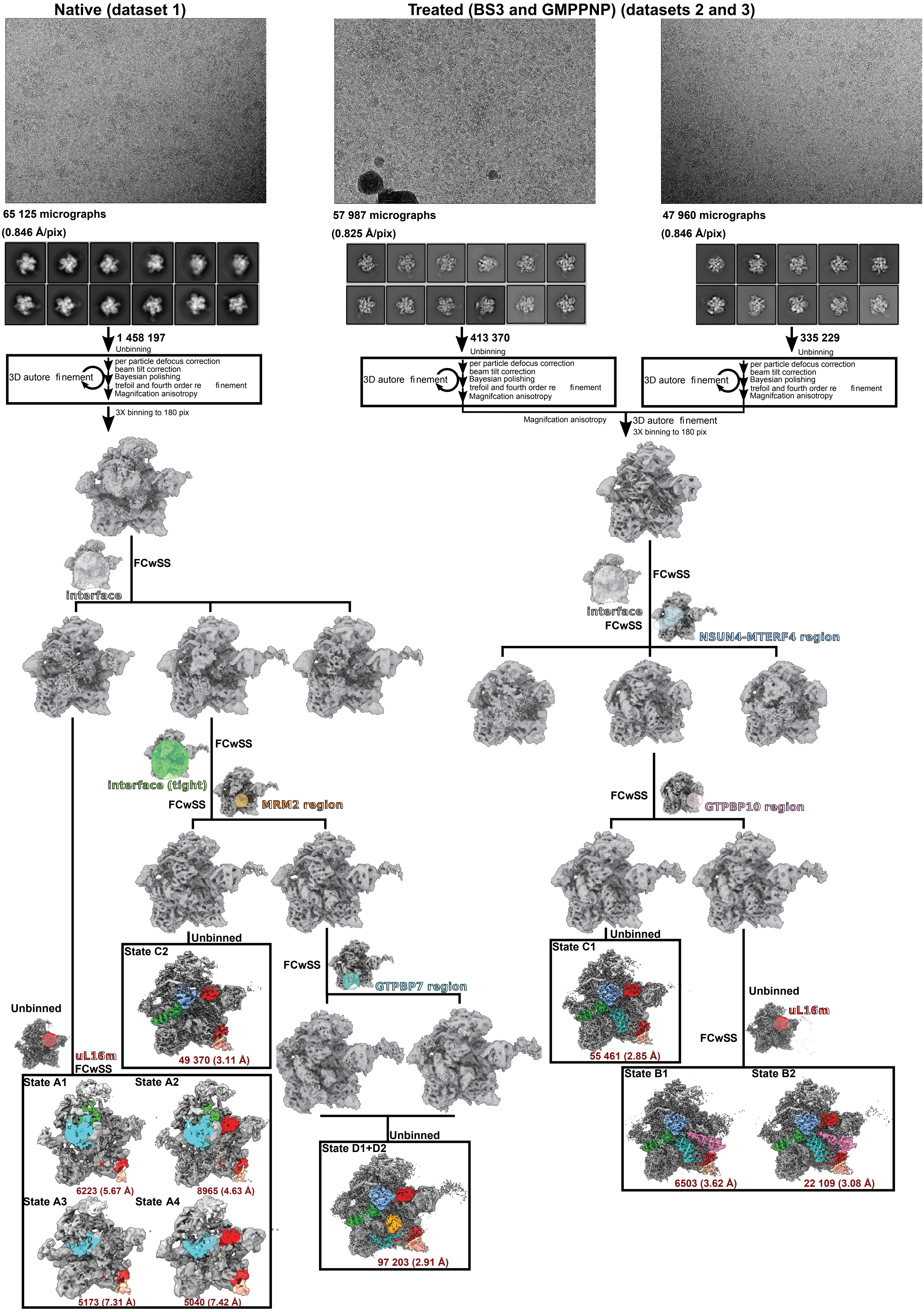


### Supplementary Fig. 10

Cryo EM data processing scheme. Top panels show representative motion-corrected micrographs followed by representative 2D classes. Remaining panels show the scheme employed for focussed 3D classification with signal subtraction (FCwSS) to resolve heterogeneity. Final unbinned maps are boxed and their respective particle numbers and resolution are indicated (red font). The resolutions are estimated using Fourier Shell Correlation (FSC) between respective half maps of 0.143 as the cut-off.

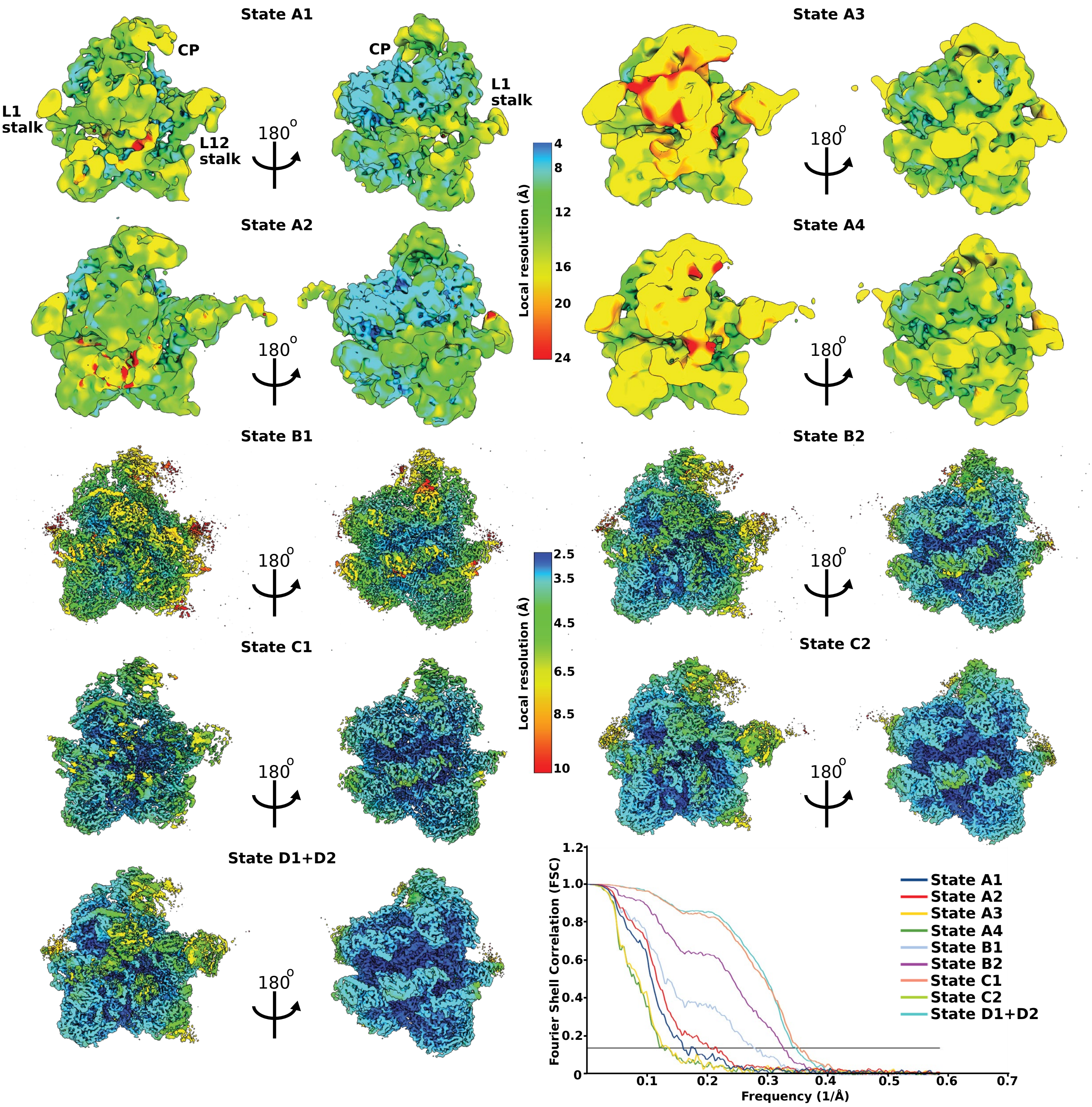


### Supplementary Fig. 11

Local resolution maps. The overall maps for all mtLSU assembly states are shown coloured by local resolution. The local resolution range is indicated by colour key. A separate range is used for states A1-A4 and B1-D2, respectively. Bottom right panel shows the FSC curves between half-maps for all the states; FSC=0.143 is marked by a black line.

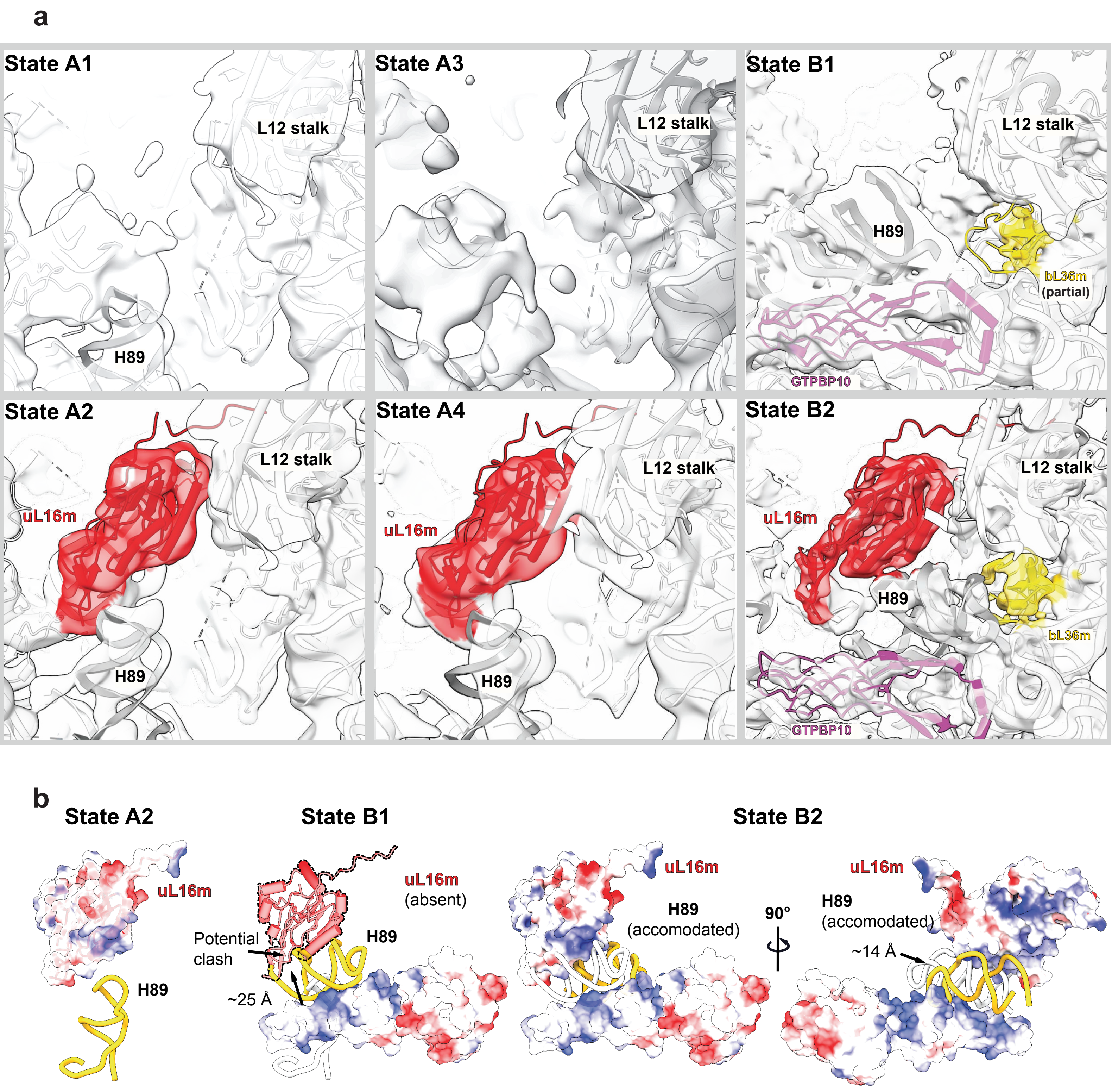


**Supplementary Fig. 12: Accommodation of uL16m, bL36m and maturation of H89.**
**a**, States A1-A4 (left and center) show unstable binding of uL16m, while, bL36m is completely absent. H89 s modeleld into a weak density in proximity to uL16m in states A2 and A4; but becomes more disordered in the absence of uL16m in states A1 and A3 . States B1 and B2 (right) feature the binding of GTPBP10. In state B1, uL16m is absent, while bL36m is observed for the first time, but, at partial occupancy as indicated by a weak density. In state B2, a stable binding of both uL16m and bL36m is observed and H89 adopts its final conformation. **b**, The panels depict the conformational rearrangement of H89 from A2 (left) to its mature state in B2 (right), guided by uL16m and GTPBP10 (colored by electrostatic potential). In each panel, the current conformation of H89 (gold) is compared with that in the previous state (white). Absence of uL16m in state B1 is highlighted by a dotted outline.
